# Genetic variation regulates the activation and specificity of Restriction-Modification systems in *Neisseria gonorrhoeae*

**DOI:** 10.1101/630434

**Authors:** Leonor Sánchez-Busó, Daniel Golparian, Julian Parkhill, Magnus Unemo, Simon R. Harris

## Abstract

Restriction-Modification systems (RMS) are one of the main mechanisms of defence against foreign DNA invasion and can have an important role in the regulation of gene expression. The obligate human pathogen *Neisseria gonorrhoeae* carries one of the highest loads of RMS in its genome; between 13 to 15 of the three main types. Previous work has described their organization in the reference genome FA1090 and has experimentally inferred the associated methylated motifs. Here, we studied the structure of RMS and target methylated motifs in 25 gonococcal strains sequenced with Single Molecule Real-Time (SMRT) technology, which provides data on DNA modification. The results showed a variable picture of active RMS in different strains, with phase variation switching the activity of Type III RMS, and both the activity and specificity of a Type I RMS. Interestingly, the Dam methylase was found in place of the NgoAXI endonuclease in two of the strains, despite being previously thought to be absent in the gonococcus. We also identified the real methylation target of NgoAX as 5’-GCAGA-3’, different from that previously described. Results from this work give further insights into the diversity and dynamics of RMS and methylation patterns in *N. gonorrhoeae*.

## Introduction

*Neisseria gonorrhoeae* is a sexually-transmitted pathogen that causes gonorrhoea. Antimicrobial resistance in this pathogen is of great public health concern and there have recently been an increasing number of publications aimed at analysing its transmission and the evolution of genetic determinants of antimicrobial resistance using high throughput short-read sequencing^1–4^. However, Single Molecule Real-Time (SMRT) PacBio sequencing provides deeper information about microbial genomes, including data on DNA modification, mainly methylation in the form of 6mA, 5mC or 4mC. Several studies have focused on the reference genome *N. gonorrhoeae* FA1090 to characterize the population of Restriction-Modification systems (RMS) in the gonococcus^5–8^ and its methylation landscape^9^. However, RMS and methylation status are known to vary significantly between strains of the same species, so this provides only limited knowledge about RMS and methylation in the gonococcus as a whole.

*N. gonorrhoeae* is among the bacterial species with the highest numbers of RMS despite being naturally competent for DNA uptake^10^. Indeed, highly transformable bacteria have been found to contain a higher number of RMS compared with those which are less competent, acting as a defence system against the invasion of foreign DNA^11^. RMS are formed by a restriction endonuclease (REase) and a DNA methyltransferase (MTase) that recognize a specific pattern in the genome. If the MTase is active, it will methylate the associated motifs and thus protect them from cleavage by the cognate REase. Bacteria can contain up to four types of RMS, although only Types I, II and III have been found in *Neisseria*^10,12^. Briefly, Type I RMS consist of three subunits named *hsdM* (MTase), *hsdS* (specificity, S, unit) and *hsdR* (REase)^13^ that form a multi-enzyme complex that recognizes specific asymmetric sequences, methylating both strands. Type II RMS are generally formed by individual MTase and REase enzymes that recognize the same short palindromic motifs and perform a two-strand methylation or cleavage, respectively. Finally, Type III RMS forms a complex of Mod (MTase) and Res (REase) subunits^14^ that recognize specific short asymmetrical motifs. The specificity of Type I RMS is encoded by an independent gene (*hsdS*) that must be active for the other subunits to be functional, while Type III RMS harbour a DNA recognition domain (DRD) within the MTase^12^. RMS have also been associated with other roles apart from defending the genome against foreign DNA invasion, such as being involved in the epigenetic control of gene expression^11^. They have been described as selfish elements, tending to propagate on mobile genetic elements and promoting their own survival^15^. They are often associated with mobility genes, such as integrases or transposases and are sometimes flanked by repeats^16^. Indeed, they have been shown to have an important role in genetic flux among bacteria, as cleavage by REases provides fragments of double-stranded DNA that could be incorporated into the host genome^11,17^.

The action of some type III RMS is known to be regulated by phase variation^10,12^. The variation in the number of repeat copies in homopolymers or short tandem repeats, mainly due to slipped-strand mispairing during DNA replication, can cause the switch between a functional, non-functional or a different version of the gene^18^. It has been hypothesized that phase variation can be used by RMS to regulate genome flux^11^ and it has been shown to be associated with a random switching in the expression of multiple genes (the so-called ‘phasevarion’, for phase-variable regulon^12,19^) in *Haemophilus influenzae*^20,21^, *N. meningitidis, N. gonorrhoeae^5^, Helicobacter pylori*^22^, and *Moraxella catarrhalis*^23^. Apart from the type III RMS, the type II ‘orphan’ Dam MTase has also been shown to be involved in gene regulation in several Gram-negative bacteria^18^ and is even required for virulence in some pathogenic organisms such as *Escherichia coli, Salmonella, Yersinia* or *Vibrio* species^24^. However, the Dam MTase is also associated with the regulation of phase variation and participates in the methyl-directed mismatch repair system. Thus, Dam-defective bacteria have more flexibility to undergo phase variation^25^. The Dam MTase^26^ has been found in some *N. lactamica* and *N. meningitidis* strains, however, there have been no reports of *N. gonorrhoeae* carrying this enzyme until now. Instead, the gonococcus and other Dam-defective *Neisseria* contain a *dam replacing gene* (*drg*), characterized as an endonuclease^26^ that has been reported to be important for adhesion and biofilm formation^25^.

In this study, we characterised the population of RMS and their associated methylation specificities in 25 *N. gonorrhoeae* strains using SMRT PacBio data. Results from this study give a more comprehensive insight into the link between genomics and epigenomics in the gonococcus.

## Results

### Detection of active RMS and associated DNA methylation

From 13 to 15 complete RMS were found in each of the 25 *N. gonorrhoeae* strains included in the study (Supplementary Table 1): 2 of Type I, 11 of Type II and 2 of Type III, all of which are present in the REBASE database (Figure 1). A detailed scan of the genomes for genes with a Pfam annotation associated with an REase or an MTase did not detect any new RMS. Most of the systems were found surrounded by core genes coding for essential functions, such as tRNA aminoacylation during protein translation, ATP binding or magnesium ion binding, but also DNA transposition and isomerase activity (Supplementary Table 2). This could potentially be related to the fact that Type I REases have an important requirement of ATP for cleavage, and Type II REases require magnesium ion as a cofactor^13^. The amino acid translation of the repetitive region in both Type III RMS is rich in prolines and, interestingly, the *ppiA* gene upstream the methylase in NgoAX is an isomerase that catalyzes the isomerization of peptide bonds in prolyl residues, assisting protein folding^27^.

**Figure 1.**
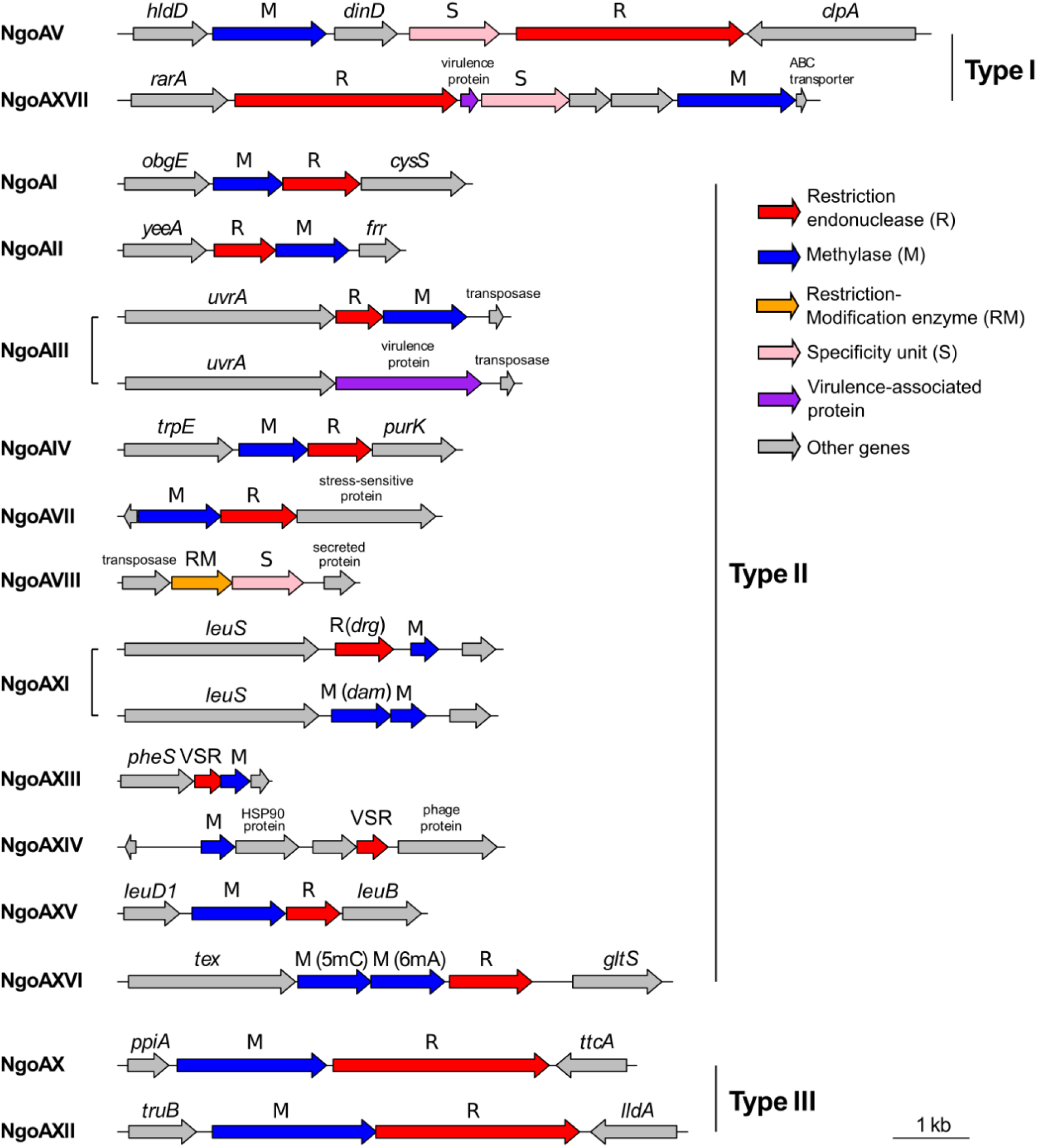
Genome organization of restriction-modification systems in *N. gonorrhoeae*. VSR: Very short patch repair endonuclease.

Analysis of patterns of DNA modification using PacBio sequencing information for the 25 gonococcal strains produced a curated list of 13 different methylated motifs (Table 1, raw motif summaries in Supplementary Table 3). Manual curation and visualization of per-base IPD ratios for each motif was essential to filter out low-frequency motifs, as several of them were found to overlap with those methylated in a higher frequency or were predicted because of higher IPD values in guanines. Indeed, O-6-methylguanine modification has been associated with DNA damage by alkylating agents^28^, and such modifications could be detected by the SMRT technology.

**Table 1.**
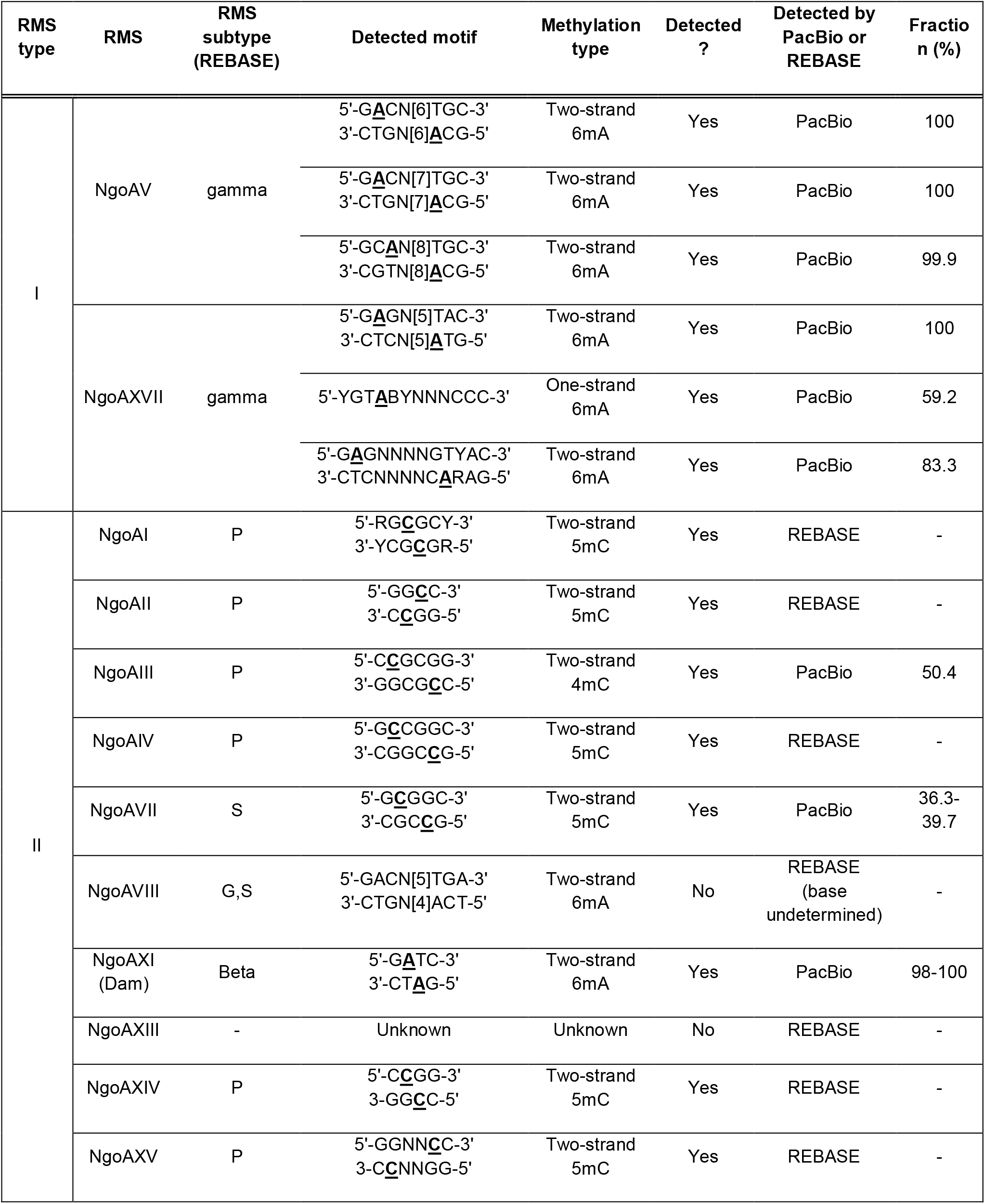
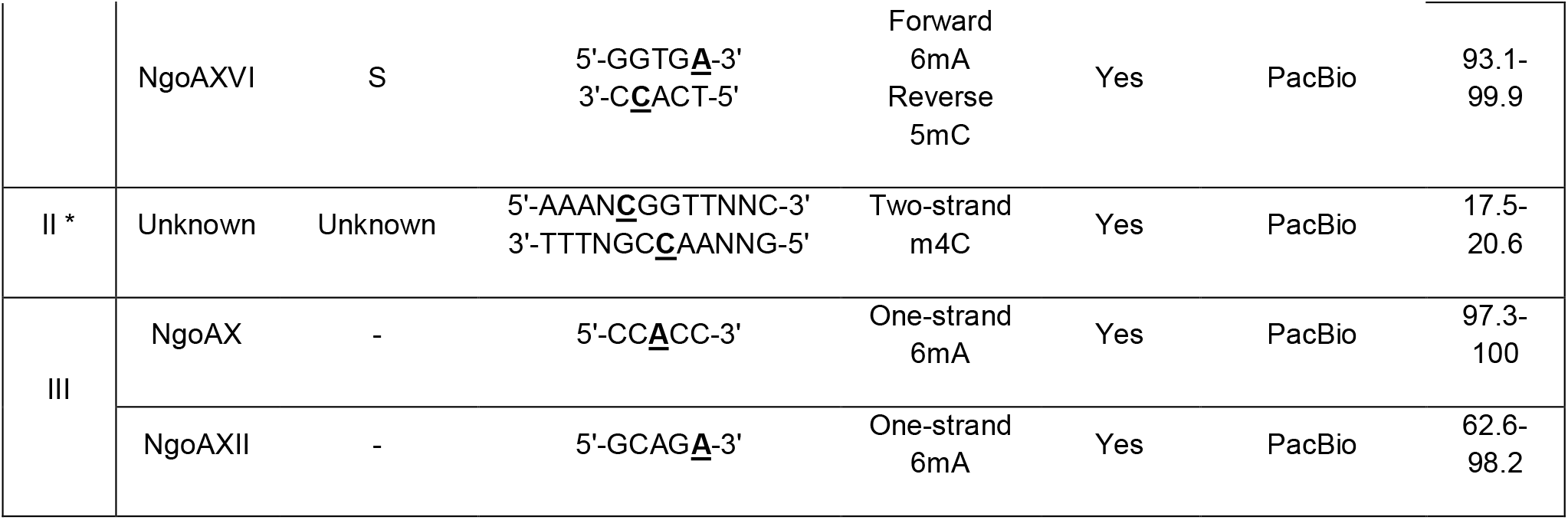
Restriction-Modifications systems and target motifs in *N. gonorrhoeae*. Those detected as methylated by the PacBio SMRT pipeline are marked as such and the fraction of modified motifs indicated. For those not directly detected by the pipeline, the associated motifs in REBASE are shown and marked as detected if a significant difference in IPD ratios is found between the target base and the unmethylated equivalent by a Mann-Whitney test.

A general investigation of the average IPD ratios for methylated and unmethylated bases was conducted. 6mA modifications extracted from our final motif list showed the highest IPD values, with a mean above 5 (quantile (Q)25 = 4.3, Q75 = 6.7; Supplementary Figure 1). Cytosine methylations (4mC and 5mC) are difficult to detect using native DNA, specially 5mC, because of the lower effect on delaying the polymerase during PacBio sequencing. Still, they showed an average IPD ratio of around 3 (Q25 = 2.3, Q75 = 3.6 for 4mC; Q25 = 2.2, Q75 = 2.9 for 5mC), higher than the mean values for any unmethylated base (Q25 = 0.9, Q75 = 1.3; Supplementary Figure 1).

### Type I RMS: NgoAV is multispecific

Two Type I RMS were present in the *N. gonorrhoeae* strains (Figures 1 and 2). Type I NgoAV RMS showed at least three different sources of genetic variability within the specificity unit (*hsdS*). An insertion of a T at position 186 of the gene creates a frameshift that causes a premature stop codon in position 62 of the protein that inactivates *hsdS*, and thus, the whole RMS (Figure 3). From 0 to 3 ATLE repeats were observed in the amino acid sequence in different strains. Some strains also contained a downstream frameshift caused by the deletion of two Gs in a homopolymer. This second frameshift does not cause the inactivation of the gene, but in combination with the number of ATLE repeats, was found to be associated with a change in the recognition pattern (Figure 3 and Supplementary Table 4). Three different methylation targets were associated with this RMS. Strains NCTC10931 and WHO M with 1 ATLE repeat and no frameshift showed two-strand 6mA methylation in the 5’-GACN[6]TGC-3’ motif (Supplementary Figure 2). However, four strains with 2 ATLE repeats instead showed the same starting and ending triplet nucleotides as the recognition pattern but the spacer was 1 bp longer, 5’-GACN[7]TGC-3’ (Supplementary Figure 3). Finally, two strains (FA1090 and WHO W) showed a slightly different motif which is a further 1 bp longer, 5’-GCAN[8]TGC-3’ (Supplementary Figure 4). In this case, they also contained 2 ATLE repeats but with the frameshift downstream that causes a premature end of the protein, which therefore does not inactivate it but changes its specificity (Figure 3). The only exception was strain NCTC12700, which contained an apparently active methylase (*hsdM*) and complete specificity unit but no associated methylated motif. This could be caused by *hsdM* or *hsdS* being not functional due to unrecognised regulatory or mutational changes.

**Figure 2.**
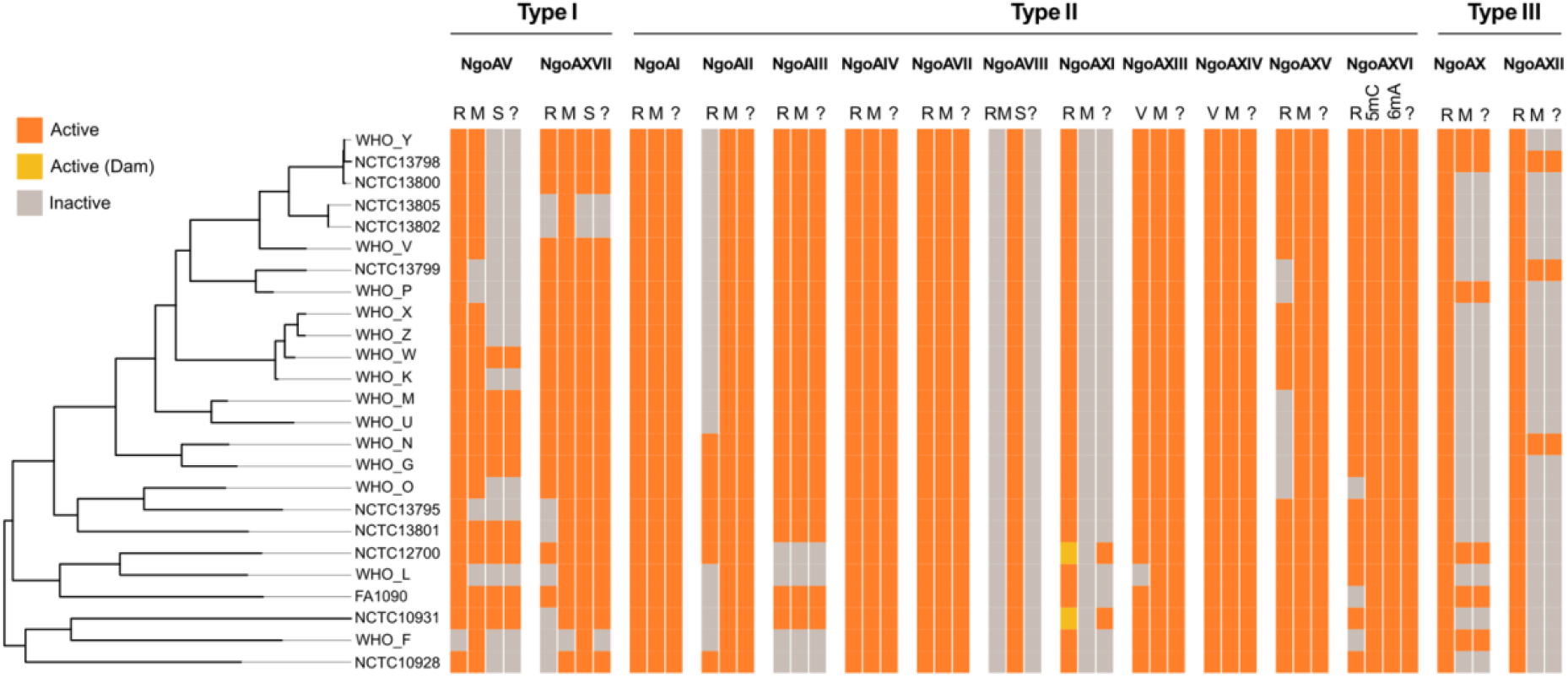
Active and inactive restriction-modification systems (RMS) in the 25 *N. gonorrhoeae* strains under study. Phandango visualization of active and inactive subunits of each RMS. The Dam methylase is marked in a different colour (see legend). An extra column (‘?’) in each RMS is plotted showing if methylation is expected for each strain. The tree on the left is a maximum likelihood reconstruction of the core genome SNPs of the 25 strains.

**Figure 3.**
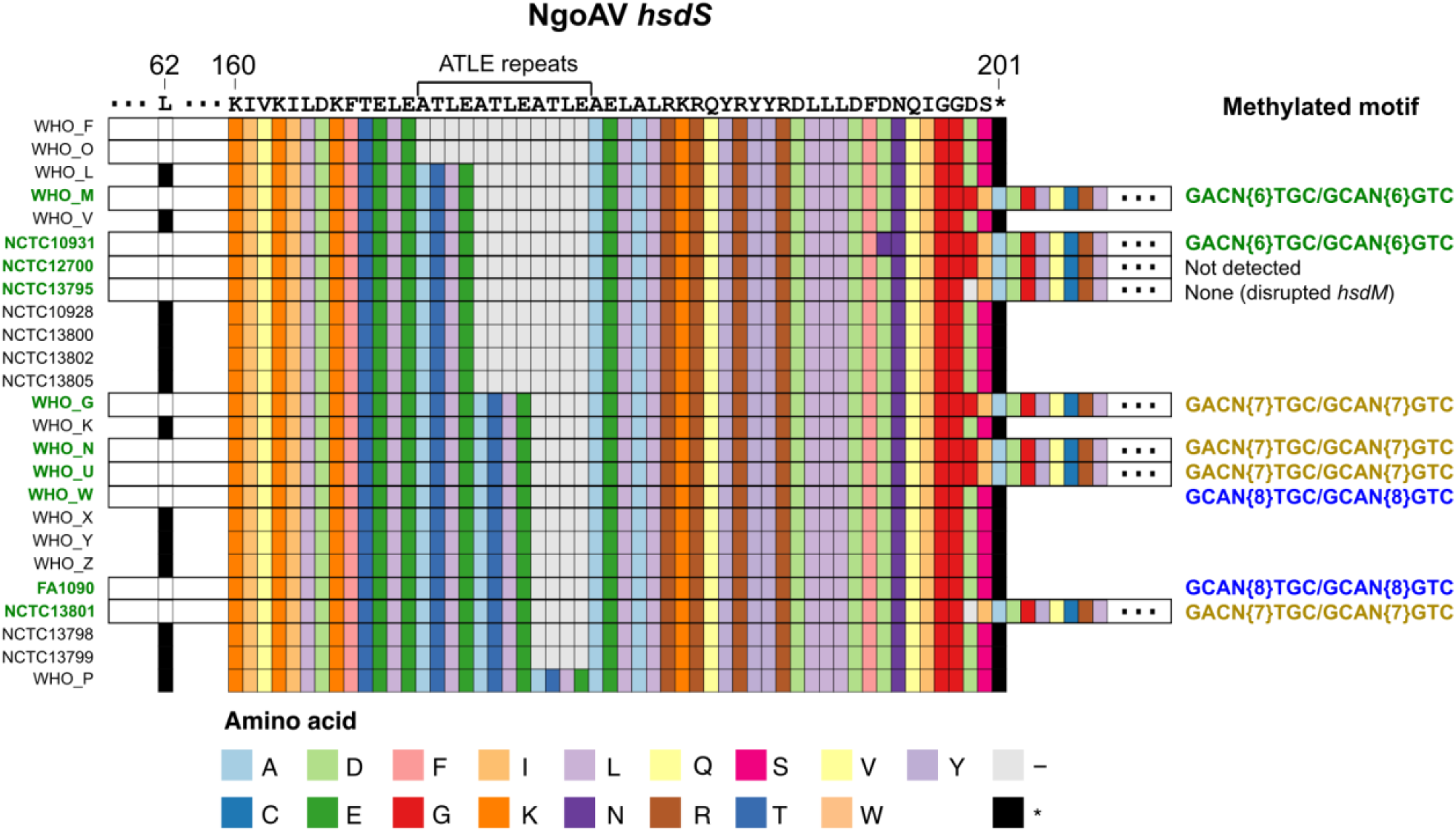
Protein alignment of the *hsdS* specificity unit of the Type I NgoAV RMS. The three main sources of variation within the unit are shown as a premature stop codon in position 62 of the protein that causes inactivation of the system, a variable number of ATLE repeats and a downstream stop codon in position 201 which, in combination, are related to a different recognition motif. The strains with a complete *hsdS* are labelled in green (left), and the detected methylated motifs using PacBio are shown in different colours on the right. ‘-’ represents gaps and ‘*’ stop codons.

The second Type I RMS, NgoAXVII, did not show specific regions of variability, only the inactivation of the methylase and/or the specificity unit causes the inactivation of the whole RMS. All strains with an active *hsdM* and *hsdS* in this RMS showed two-strand 6mA methylation in the 5’-GAGN[5]TAC-3’ motif (Supplementary Figure 5 and Supplementary Table 5). Two additional motifs were detected in three strains (FA1090, NCTC12700 and WHO O) that we hypothesized are off-target methylations by this enzyme because of the similarity of the motifs: double-stranded methylation in 5’-GAGN[4]GTYAC-3’/3’-CTCN[4]CARTG-5’ and single-stranded methylation in 5’-YGTABYN[3]CCC-3’, which is the reverse complement of 5’-GGGN[5]TAC-3’, very similar to the main motif methylated by NgoAXVII, 5’-GAGN[5]TAC-3’ (Table 1). The specificity unit of this enzyme was found to be very conserved, with only two non-synonymous mutations that had no association with the observed off-target methylations.

### Cytosine methylation by Type II RMS

*N. gonorrhoeae* contains up to 11 Type II RMS (Figures 1 and 2), 7 of them strictly associated with cytosine methylation (mC). In spite of apparently showing functional cytosine methylases, only three motifs containing 5mC or 4mC methylation were detected by the PacBio pipeline (Table 1 and Supplementary Table 3). However, as we show above, both types of cytosine modification can be distinguished from an unmethylated base (Supplementary Figure 1). In order to perform a deeper analysis of the motifs associated with mC in Type II RMS, we specifically identified those reported in REBASE and evaluated their per-base IPD ratios (Supplementary Table 6). A Mann-Whitney test was performed between the distribution of IPD values for the predicted mCs and a random set of unmethylated cytosines for each strain to assess whether there was a statistically significant difference. Significant cytosine modification was found for the motifs associated with the 7 RMS expressing an active methylase (Supplementary Figure 6 and Supplementary Table 6). Four strains showed a gene annotated as “virulence-associated” instead of the NgoAIII RMS (WHO F, WHO L, NCTC10928 and NCTC12700) and three of those did not show significant signals of methylation (Supplementary Table 6). The motif associated with NgoAIII in REBASE is 5’-GGCGCC-3’ (5mC) and this was detected as a 4mC signal (reported as the degenerate motif 5’-GG<u>C</u>SCCND-3’) in WHO V (Supplementary Table 3), although we showed the signal was in fact present in all the strains with a functional RMS (Supplementary Figure 6). The WHO G data produced a non-significant test for mC in the forward strand of 5’-G<u>C</u>GGC-3’, associated with NgoAVII, which could be due to hemi-methylation or poor signal in the data.

RMS NgoAXIII was found as a VSR (Very Short patch Repair) protein followed by a MTase and there is no report of its target motif in REBASE. A series of BLAST searches revealed that only *N. polysaccharea* M18661 (Genbank accession number CP031325.1) contained a homologous region to NgoAXIII. Instead, genomes of *N. lactamica* (i.e. Y92-1009, Genbank accession number CP019894.1) showed a longer version of the MTase (362 amino acids compared to 112 in *N. gonorrhoeae* FA1090 and 133 in the rest). This finding could mean that *N. gonorrhoeae* does not have a complete NgoAXIII system, but only remnants of a non-active MTase. Interestingly, several *N. meningitidis* genomes contain the VSR protein followed by a different RMS instead (NmeDI).

The 5’-AAA<u>C</u>GGTTNNC-3’ motif was detected by the PacBio SMRT pipeline containing single-strand m4C methylation in the underlined base. The study of the distribution of per-base IPD ratios in the WHO strains revealed that the complementary sequence was probably also methylated at 5’-AAANCGGTTNNC-3’/3’-TTTNGC<u>C</u>AANNG-5’ (Supplementary Figure 7). A Mann-Whitney test revealed the distribution of IPD values for these two bases to be significantly different than that of an unmethylated cytosine for most of the strains (Supplementary Figure 8). FA1090 and, especially, the NCTC strains, consistently showed very noisy results, but the test was still statistically significant for them (Supplementary Figure 8). We hypothesize that the methylation of this motif could result from a secondary target of a Type II RMS or might be the target of NgoAXIII, for which the methylated motif is currently unknown, although this is unlikely as the fraction of methylated 5’-AAANCGGTTNNC-3’ is very low (Table 1).

### 6mA methylation by Type II RMS

Only three of the Type II RMS contain 6mA methylases. The NgoAVIII system has a BcgI-like structure^29^ as it is formed by an enzyme with restriction and 6mA methylation properties (RM) and a specificity unit (S). A BLASTn search of the genomic region spanning both units against the NCBI database revealed that this RMS has not been described in any other organism. A transposase immediately upstream of the RM unit indicates that this system may be mobile and acquired from an unknown source. The target motif in REBASE for this RMS is that of BcgI, 5’-GACN[5]TGA-3’, although we do not detect this motif as methylated or any other motif with 6mA methylation that could be associated with NgoAVIII (Supplementary Table 3), despite the system appearing intact. Interestingly, BcgI-like MTases are known to prefer hemi-methylated DNA, although their recognition sequences are very similar to those of Type I MTases, which methylate both strands^30,31^. Thus, NgoAVIII may have a maintenance role for the motifs targeted by Type I MTases, methylating the growing strand immediately after chromosomal replication, a point when these motifs will be hemi-methylated.

The Dam methylase was found in place of the restriction enzyme in NgoAXI in two strains (NCTC10931 and NCTC12700), which showed double-strand 6mA 5’-GATC-3’ methylation (Supplementary Figure 9). This MTase has been reported to be substituted by the Dam-replacing endonuclease (Drg) in many *Neisseria* species, but has not been found in *N. gonorrhoeae* until now^25,26^. Finally, the NgoAXVI RMS is formed by two methylases (5mC and 6mA, respectively) and a restriction enzyme. A clear 6mA signal is found in the associated 5’-GGTG<u>A</u>-3’ motif in all the strains (Supplementary Figure 10). A more detailed comparative analysis of the distribution of IPD values of the three cytosines in the reverse complement of this motif against unmethylated Cs for each strain identified the middle cytosine as the most plausible candidate for 5mC methylation (3’-C<u>C</u>ACT-5’ Supplementary Figure 11).

### Type III RMS: the real target of NgoAXII is 5’-GCAGA-3’

Two Type III RMS were present in the *N. gonorrhoeae* strains (Figures 1 and 2). These contain a phase-variable methylase (*Mod*) that controls the activation or inactivation of the system and includes a DNA-recognition domain (DRD) that controls its specificity. NgoAX contained a variable number of 5’-CCCAA-3’ repeats that switch the system ON/OFF, and the *modB1* DRD allele (Supplementary Table 7). In contrast, the NgoAXII contained a variable number of 5’-AGCC-3’ repeats and the *modA13* DRD allele (Supplementary Table 8). Both *modA* and *modB* DRD alleles are those typically found in *N. gonorrhoeae*^8,32^.

Active NgoAX RMS showed 6mA 5’-CC<u>A</u>CC-3’ methylation in 6 strains (Supplementary Figure 12 and Supplementary Table 7), and active NgoAXII showed 6mA 5’-GCAG<u>A</u>-3’ methylation in 3 strains (Supplementary Figure 13 and Supplementary Table 8). Interestingly, a previous report suggested the methylated motif for the *modA13* allele of this enzyme was 5’-AGA<u>A</u>A-3’ by evaluating the digestion pattern of ApoI and other enzymes^5^, and this is the recognition motif associated with NgoAXII in REBASE. However, our analysis of the distribution of the per-base IPD ratios for both motifs in the three strains with an active *modA* (NCTC13798, NCTC13799 and WHO N; Figure 4) revealed that less than 0.1% of the AGA<u>A</u>A motifs contained an IPD ratio above 3, while most of the GCAG<u>A</u> (>79%) showed high IPDs in the underlined base (Supplementary Table 9). We only observed between 4 and 8 motifs in the genomes that included both motifs overlapping with each other and that expanded to include a complete ApoI recognition sequence (5’-RAATTY-3’), 5’-GCAG<u>**A**AATTY</u>-3’. Therefore, the digestion in these cases could have been impeded by the methylation of the second A instead (in bold). A low fraction of motifs with high IPDs in the first A of 5’-GC<u>A</u>GA-3’ was found in the strains with an active Type I NgoAV, and this may be due to overlapping signals between both RMS (Supplementary Figures 1-3 and Supplementary Figure 13). Strangely, results from the PacBio motif analysis in FA1090 for 5’-GCAGA-3’ did not show signals of methylation, although Mod was apparently complete in the reference genome. An assembly of the PacBio reads associated with the methylation data for this strain^9^ revealed that the methylase in the NgoAXII RM system in the subclone of FA1090 used for sequencing in the cited work contained 35 AGCC repeats instead of 37 in the reference genome, causing a protein truncation and thus a lack of 5’-GCAG<u>A</u>-3’ methylation.

**Figure 4.**
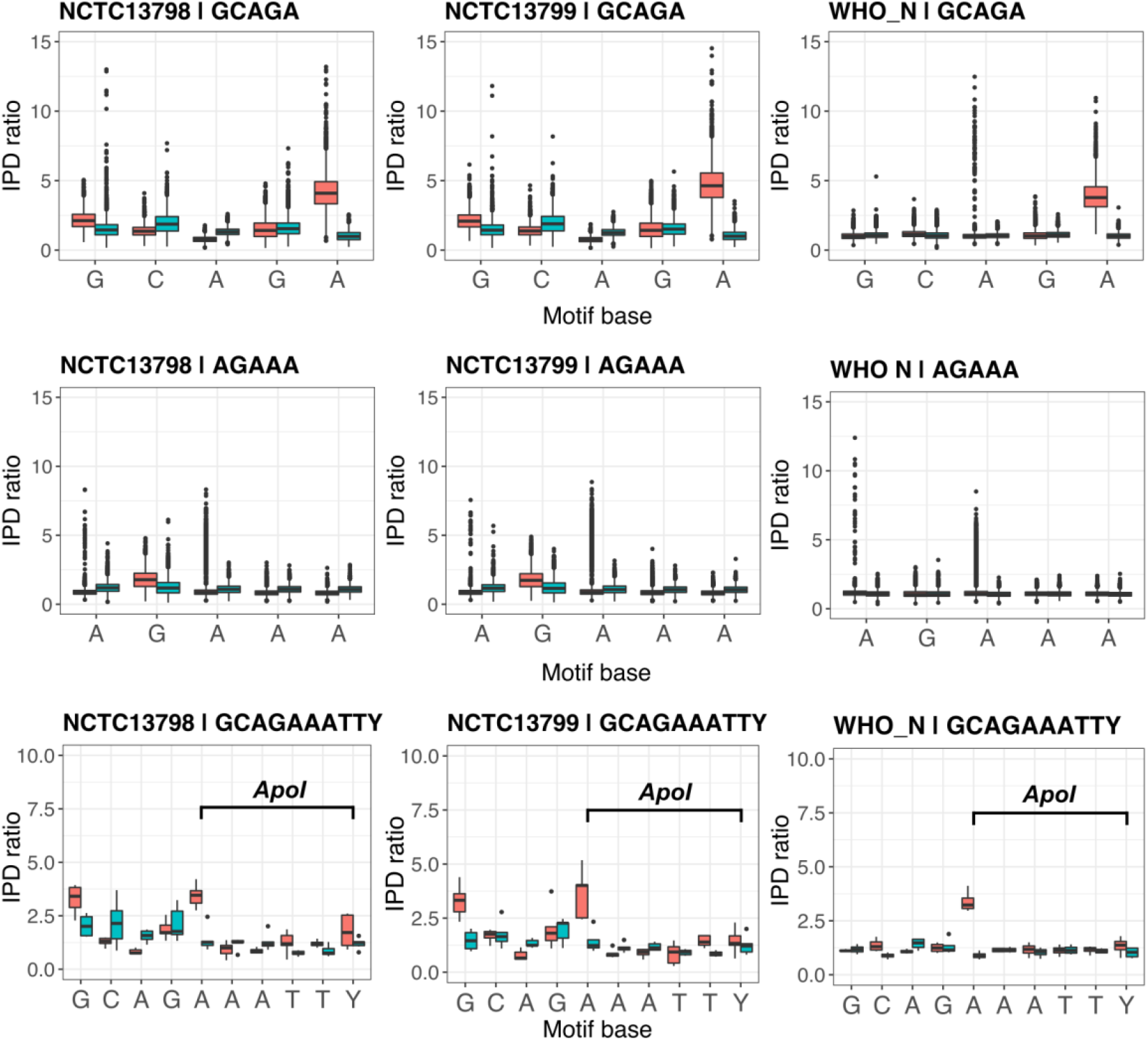
Comparison of the per-base distribution of IPD ratios for the 5’-GCAGA-3’ motif detected in this study as associated with NgoAXII, the previously inferred 5’-AGAAA-3’ and the motif overlapping among the previous two and the recognition pattern of the ApoI restriction enzyme, 5’-RAATTY-3’ (5) in the three strains with a functional NgoAXII system. Two boxplots are shown per base, corresponding to the IPD ratios of the forward (red) and reverse (blue) strands.

### Orphan methylases in *N. gonorrhoeae* include the Dam methylase

Disrupted methylases can cause the inactivation of the whole RMS, due to mechanisms that avoid self-degradation of the chromosomal DNA^12^ in this situation. However, methylases can be functional with a disrupted or absent endonuclease. These methylases have been associated with a regulatory rather than defensive role and have been described as ‘orphan’. In the strains under study we observed multiple examples of this in the Type I NgoAXVII (Supplementary Table 5) and several Type II RMS (Supplementary Table 6). The Dam methylase in NgoAXI acts as an ‘orphan’ methylase, as it completely replaces the endonuclease of the system, and the downstream MTase is disrupted. A BLASTp search of the NCBI non-redundant protein database revealed that the gonococcal Dam sequence has 97-98% amino acid identity to *N. polysaccharea*, 97% to *N. flavescens*, 96-97% to *N. lactamica*, and 92-96% to *N. meningitidis*. A broader screening of the Dam protein in the *Neisseria* genus showed a reasonable amount of diversity at the amino acid level, with recombination swapping sequence variants among *Neisseria* species (Supplementary Figure 14).

A scan of the Pfam domains of the proteome of the 25 strains under study did not reveal further DNA methylases without an accompanying restriction enzyme in the main chromosomes. However, the strains carrying the conjugative plasmid (WHO G, L, M, N, O, W, and NCTC10931) showed an orphan MTase in this element that has a 100% amino acid and nucleotide identity to the Type II M.Ngo8107ORF11P and M.Ngo5289ORFAP enzymes in REBASE. Visualization of the annotation of the WHO strains revealed that it is present in the conjugative plasmid (Supplementary Figure 15). No further matches were found to any other bacteria in a BLASTn search against a non-redundant nucleotide database.

## Discussion

Restriction-modification systems have been shown to perform several other roles apart from being a defence system against foreign DNA^11^. For example, some of them, especially those from type III, are involved in the control of gene expression through phase variable repeats in the Mod unit^19^. Previous papers have characterised different RMS in the *N. gonorrhoeae* FA1090 reference strain^6–8,33^ and also data on its methylation status has been described^9^. In this manuscript, we provide a detailed view of the structure and variability of DNA MTases and specificity units in RMS in 25 strains of *N. gonorrhoeae* and link them to the detected methylated motifs.

The analysed gonococcal genomes showed from 13 to 15 complete RMS, all of them described in the reference strain FA1090 in REBASE^34^. However, a detailed screening of the genome assemblies did not reveal any new systems or orphan MTases, apart from an enzyme homologous to M.Ngo8107ORF11P on the conjugative plasmid in the strains carrying this element. Thirteen curated methylated motifs were detected in the 25 strains, most of them containing 6mA methylation (Table 1). A detailed analysis of the genetic variability of the *hsdS* units of Type I MTases and the phase variable repeats in the DRD of Type III MTases allowed us to link them to particular methylated motifs. In the case of the Type I NgoAV, earlier work confirmed the functionality of the naturally truncated *hsdS* in FA1090, which was predicted to recognize the 5’-GCAN[8]TGC-3’ motif^35^. Later, it was proven that the cause of this truncation was a phase-variable G homopolymer and that the restitution of a complete form of this unit changed the specificity of the MTase to 5’-GCAN[7]STGC-3’^7^. Our study reveals further insight into the diversity of specificities of the NgoAV RMS. Three different motifs were detected in strains with a functional NgoAV. The change in specificity was found to be related to a combination of two factors: the number of ATLE repeats in the protein sequence of *hsdS* and the presence of the frameshift that causes the known truncation downstream of these repeats (Figure 3 and Supplementary Table 4). Strains with 1 ATLE repeat and a complete *hsdS* showed 5’-GACN[6]TGC-3’ double-stranded 6mA methylation, strains with 2 ATLE repeats and a complete *hsdS* showed 5’-GACN[7]TGC-3’ and strains with 2 ATLE repeats and a downstream frameshift causing truncation, which is the case in FA1090, showed 5’-GCAN[8]TGC-3’ (Supplementary Table 4) instead. The presence of amino acid repeats in the centre region of *hsdS* has already been described^35,36^, but not its role in sequence specificity in combination with the downstream frameshift. Off-target methylated motifs were associated with the second Type I RMS NgoAXVII, although no associated variability was found in *hsdS* in this system.

Seven out of 11 Type II RMS in *N. gonorrhoeae* are associated with cytosine methylation. However, this type of DNA modification is difficult to detect by current PacBio sequencing, and high coverage in combination with Tet-conversion is preferable^37^. In this study, we relied on available data from other projects, which performed sequencing on untreated native DNA, thus, most of the analysis were performed on 6mA methylation. Only three motifs associated with 5mC or 4mC methylation were detected in a subset of the strains (Table 1). Nonetheless, a statistical comparison between the IPD values of target cytosines in the motifs annotated in REBASE for these RMS to those from unmethylated cytosines revealed significantly higher values for those in the strains carrying active 5mC MTases (Supplementary Table 5). 5mC can be converted into T by deamination, generating mismatches that can produce mutations. Bacteria can modulate this effect by methylating 4mC instead^11^. In fact, double-stranded methylation of 5’-CCGCGG-3’ by NgoAIII was detected by the PacBio system as 4mC, apart from the potential off-target 5’-AAANCGGTTNNC-3’ motif (Table 1). Interestingly, secondary methylation by these types of enzymes has been observed in *H. influenzae*^38^.

*N. gonorrhoeae* harbours two VSR endonucleases to correct T:G mismatches, V.NgoAXIII and V.NgoAXIV. The specificity of NgoAXIII is still unknown and, in this study, we hypothesize that the MTase may be truncated, as a longer version is found in *N. lactamica* (Genbank accession number CP019894.1). However, the VSR may still be functional, as it has been found to recognize T:G mismatches in every nucleotide context^33^. V.NgoAXIV has also been described to recognize mismatches in sequences other than 5’-CCGG-3’^33^. No methylated motif has been found for the Bcg-like RMS NgoAVIII, however, previous reports describe this type of MTase as having a strict preference for hemi-methylated sites^30,31^ in Type I-like motifs. Thus, we propose that NgoAVIII may have a maintenance role, methylating the replicated strand in sites targeted by Type I RMS during DNA replication.

The Dam methylase is present in some strains of *N. lactamica* and *N. meningitidis*^26^. However, it has not been described in *N. gonorrhoeae* until now. Instead, *Neisseria* species or strains lacking this enzyme are known to carry an endonuclease encoded by *drg* (*dam* replacing gene) recognizing the 5’-GATC-3’ motif^26,39^. Here, we observed two *N. gonorrhoeae* strains (NCTC10931 and NCTC12700) carrying the Dam MTase, instead of the *drg* gene, in the NgoAXI RMS locus, followed by a truncated MTase. Thus, Dam is acting as an orphan enzyme, a form that has been related to gene expression regulation^40^. Interestingly, previous publications have suggested that strains carrying the *drg* gene have an advantage over those carrying the Dam MTase because they have more flexibility for phase-variation, as Dam participates in DNA mismatch repair during replication^25,39^. Also, the same publications showed experimentally that strains carrying *drg* form more stable biofilms and have better adhesion to human cells during infection.

Several works have studied the so-called ‘phasevarion’ in *N. gonorrhoeae* and *N. meningitidis*, referred to the set of genes with an altered expression due to phase variation in an MTase^5,12^. Type I NgoAV has been considered to regulate a phasevarion as the phase variation of the G homopolymer is involved in a change in specificity^19^. In our work, we complement this result, as we show that the change in specificity is produced as a combination of the phase-variable poly-G and the number of ATLE repeats in the amino acid sequence (Supplementary Table 4). However, the best example of this is the phase-variable Type III RMS^12^, in which for example *modD1* has been associated with hypervirulent *N. meningitidis* clonal complexes and *modA13* with enhanced biofilm formation and intracellular survival^19^. In a recent publication, the NgoAX Mod was experimentally inactivated in *N. gonorrhoeae* and a deregulation of the expression of 121 genes was observed, along with an effect on the adherence to and invasion of host epithelial cells, which are essential for infection^8^. In contrast, inactivation of the NgoAXII Mod enzyme showed a deregulation of 54 genes under iron-limiting conditions^5^. Studies have reported that *N. gonorrhoeae* carries a very conserved *modB1* allele in NgoAX compared to *modA13* in NgoAXII. These results have led to the suggestion that NgoAX Mod is the main regulator of the epigenome, while NgoAXII regulates in specific growth conditions^8^. Here, we observe that indeed all the strains under study carry the *modA13* and *modB1* alleles, with all possible combinations of activity apparent: we detected strains with only the NgoAX Mod active (i.e. FA1090 or WHO F), some with only the NgoAXII Mod active (WHO N and NCTC13799), one with both active (NCTC13798), and the rest with both systems inactive (Supplementary Table 7 and Supplementary Table 8).

In summary, *N. gonorrhoeae* contains several RMS to protect the genome against foreign DNA invasion. Genetic diversity created through phase variation or hypervariable domains controls their activation or inactivation, along with a variation of specificity in one case. This work gives further insight into the RMS in *N. gonorrhoeae* and the consequent methylation landscape, which is able to change to modulate gene expression. We also show the importance of detailed PacBio SMRT analysis to enhance and complete the methylation information contained in public databases.

## Methods

### Genomes included in the study

We analysed the genomes of a total of 25 *N. gonorrhoeae* strains that had been sequenced using PacBio. Of those, 14 complete genomes were obtained from the 2016 WHO reference panel^41^ and 10 additional genomes were selected from the Public Health England NCTC 3000 collection (http://www.sanger.ac.uk/resources/downloads/bacteria/nctc/) avoiding overlap with the WHO panel, to further improve the representativeness of the *N. gonorrhoeae* genome diversity (Supplementary Table 1). Additionally, PacBio raw data and predicted motifs were retrieved for the reference genome *N. gonorrhoeae* FA1090 from a recent study^9^. Sequencing was run using native DNA in all cases. Assembly and annotation were performed as reported in the publications cited above, using an automatic and improved pipeline at the Wellcome Sanger Institute^41^. FA1090 was reassembled using the PacBio data with Canu v1.6^42^ to compare the number of tandem repeats in the type III MTases with those from the original assembly available in the public databases (GenBank accession AE004969).

### Analysis of DNA methylation

Genome-wide base modifications and predicted modified motifs were called using the RS_Modification_and_Motif_Analysis protocol from the PacBio SMRT Analysis software v2.3.0. Coordinates of the predicted motifs were localized in all the genomes and plasmids using the EMBOSS application *fuzznuc*^43^. Per-base IPD ratios for the predicted motifs were extracted from the raw data and visualized in the 25 strains using R^44^.

The distribution of IPD values for each of the four bases outside any of the predicted motifs was used as the distribution of values for the four unmethylated bases. A random subsample of 10,000 unmethylated sites of each base was used. Cytosine methylation from type II RMS was inferred by evaluating the distribution of IPD ratios in the target base in the associated motif in REBASE^34^ compared to that of unmethylated cytosines. A Mann-Whitney test was performed to evaluate statistical significance and p-values were corrected using Bonferroni correction^45^.

### Detection and specificity of RMS

An extended and manually curated list of Pfam domains associated with REases and MTases from a previous work^46^ was used to detect these genes in the annotations of the genomes under study. HMMER^47^ *hmmscan* was run independently on the proteome of each genome (.faa files) against Pfam(A) v30^48^ to complement the annotation information. R language^44^ was used to parse the annotation files and the results from *hmmscan* and extract genes with an inferred Pfam domain in the list of target REases and MTases. Genes predicted to be an REase or a MTase that were less than 10 genes distant were considered to be part of the same RMS. The rest were tagged as ‘orphan’. Results were compared to the RMS annotated for FA1090 in REBASE^34^. The presence or absence of every RMS in each strain was visualized using phandango^49^.

The motif annotated in REBASE^34^ for each MTase was compared to the list of predicted motifs obtained by the PacBio SMRT Analysis software for all the strains. Nucleotide and protein sequences of every unit of the RMS were extracted for each strain and compared using SeaView v4.6.1^50^ to look for sources of variability within the MTases or the S units. These and the flanking genes were visualized using Artemis v16.0.0^51^. Disrupted proteins were confirmed by a protein-protein BLAST against a non-redundant protein database^52^.

Representative protein sequences of the Dam methylase in the *Neisseria* genus were downloaded from the Identical Protein Groups (IPG) tool of the NCBI database (accessed on 21/01/2019). These sequences were aligned using SeaView v4.6.1^50^ and the resulting alignment trimmed using Gblocks v0.91b^53^ considering only positions in which at least half of the sequences do not have a gap. PhyML^54^ was used to build a maximum likelihood tree using the LG model and performing 100 bootstrap replicates. Final tree and metadata were visualised using iToL^55^.

### GO enrichment analysis

A Gene Ontology (GO) enrichment analysis was performed for the genes flanking all the RMS using the *topGO* R package^56^. The three ontologies were scanned (‘BP’, biological process; ‘MF’, molecular function; and ‘CC’, cellular component). The *classic* and *weight01* algorithms were used which do not, and do, use hierarchy information for scoring a particular GO term, respectively^57^. The statistical significance of the enrichment was calculated using a Fisher’s exact test between the observed and expected number of genes in each term for each algorithm. Terms with a p-value<0.05 in both algorithms were considered as significant in the results. P-values were not corrected for multiple testing to avoid excluding significant GO terms near the cut-off. Besides, the tests are not independent when using the *weight01* algorithm as it is conditioned on neighbouring terms^56^.

### Data availability

Scripts used to perform the analyses and plots in this work are available in the GitHub repository https://github.com/leosanbu/MethylationProject. The 14 *N. gonorrhoeae* PacBio raw data and complete genomes from the 2016 WHO panel are available under the ENA Bioproject PRJEB14020 (Sample accessions SAMEA2448460-SAMEA2448470 and SAMEA2796326-SAMEA2796328)^41^. Accession numbers for the PacBio data from the ten strains downloaded from the NCTC3000 project are available in the following link: https://www.sanger.ac.uk/resources/downloads/bacteria/nctc/ (Sample accessions SAMEA3174297-SAMEA3174299, SAMEA4076737, SAMEA4076741, SAMEA4076765, SAMEA4076768-SAMEA4076770, SAMEA4076773). GFF files for the WHO and NCTC sequence data are available in the GitHub repository https://github.com/leosanbu/MethylationProject. The *N. gonorrhoeae* FA1090 reference genome was obtained from NCBI accession number AE004969 and the PacBio raw data from the study by Blow *et al* (2016)^9^. Supplementary Table 1 contains the detailed information for each strain.

---

Supplementary Table 1. List of *N. gonorrhoeae* strains used in the analysis. The ENA accession information of the raw and complete genomic data are indicated along with the average per-base sequencing coverage. Sample accession numbers are used for raw data as some of the strains include more than one sequencing run.

Supplementary Table 2. Results from a GO enrichment analysis of the flanking genes of the 15 RMs in *N. gonorrhoeae*. The number of genes annotated in each GO term and the expected number of genes to fall in each category are shown together with the number of significant hits in the flanking genes. GO terms significantly enriched by both the *classic* and *weight01* algorithms (p-value < 0.05) are shown. Results are shown for the three sub-ontologies (BP = Biological Process, MF = Molecular Function, CC = Cellular Component).

Supplementary Table 3. Compilation of the motif_summary.csv output files of the PacBio methylation pipeline run on the 25 N. gonorrhoeae included in the study. Fields are explained in the following technical note: https://github.com/PacificBiosciences/Bioinformatics-Training/wiki/Methylome-Analysis-Technical-Note.

Supplementary Table 4. Type I NgoAV restriction-modifications system (RMS). Active (A) or disrupted (D) components are indicated. Red highlight is used to mark a cause of disruption. The different sources of variability in the pattern recognition domain of the specificity unit, which cause different methylated motifs (in different colours) are shown. An active methyltransferase requires both the methylase and the specificity unit to be active. In that case, if the restriction enzyme is not functional, the methylase is tagged as ‘orphan’. *hsdR:* Restriction endonuclease; *hsdM*: Methyltransferase; *hsdS*: Specificity unit.

Supplementary Table 5. Type I NgoAXVII restriction-modifications system (RMS). Active (A) or disrupted (D, in red) components are indicated. An active methyltransferase requires both the methylase and the specificity unit to be active. In that case, if the restriction enzyme is not functional, the methylase is tagged as ‘orphan’ (blue). *hsdR*: Restriction endonuclease; *hsdM*: Methyltransferase; *hsdS*: Specificity unit.

Supplementary Table 6. Type II restriction-modifications systems (RMSs). Active (A) or disrupted (D, in red) components are indicated. An active methyltransferase requires both the methylase and the specificity unit to be active. In that case, if the restriction enzyme is not functional, the methylase is tagged as ‘orphan’ (blue). R: Restriction enzyme; M: Methyltransferase. The Bonferroni-corrected p-values of a Mann-Whitney test between the IPD values of methylated cytosines of each motif (as indicated in REBASE) and the distribution of unmethylated cytosines for each strain are shown. ****p-value < 0.001; ***p-value<0.001; **p-value<0.01;*p-value<0.05; ns: non-significant.

Supplementary Table 7. Type III NgoAX RM system. Active (A) or disrupted (D, in red) components are indicated. The number of CCCAA repeats and the DNA recognition domain (DRD) allele are shown, which are the source of variability in the Mod unit. An active methyltransferase requires both the methylase and the specificity unit to be active. In that case, if the restriction enzyme is not functional, the methylase is tagged as ‘orphan’. Res = Restriction enzyme, Mod = Methyltransferase.

Supplementary Table 8. Type III NgoAXII RM system. Active (A) or disrupted (D, in red) components are indicated. The number of AGCC repeats and the DNA-recognition domain (DRD) allele are shown, which are the source of variability in the Mod unit. An active methyltransferase requires both the methylase and the specificity unit to be active. In that case, if the restriction enzyme is not functional, the methylase is tagged as ‘orphan’. Res = Restriction enzyme, Mod = Methyltransferase.

Supplementary Table 9. Number of GCAGA and AGAAA and GCAGAAATTY motifs in each strain and number and proportion of those with an IPD>3 in the following underlined bases. GCAGA is the motif associated to NgoAXII detected by the PacBio SMRT analysis pipeline (NCTC13798, NCTC13799 and WHO N), AGAAA is the predicted motif for this methylase in REBASE and GCAGAAATTY is the motif resulting from overlapping GCAGA, AGAAA and the recognition sequence of the ApoI enzyme (5’-RAATTY-3’).

Supplementary Figure 1. Distribution of the IPD ratios for methylated and unmethylated bases. 6mA, 4mC and 5mC IPD values were extracted from the target bases in the final list of curated motifs. Those below an IPD ratio of 2 where excluded to minimize the inclusion of unmethylated motifs. Values for each unmethylated base correspond to a random sample of 10,000 sites per strain outside all the non-redundant predicted motifs detected by the PacBio SMRT pipeline. Dashed vertical lines and numbers indicate the mean value of each distribution.

Supplementary Figure 2. IPD ratio values for each base in all instances of the 5’-GACN[6]TGC-3’ motif, target of the Type I NgoAV RMS, in each of the 25 *N. gonorrhoeae* strains included in the study. Per-base distribution of IPD ratio is shown as two separate boxplots containing the values in the forward (red) and reverse (blue) strand.

Supplementary Figure 3. IPD ratio values for each base in all instances of the 5’-GACN[7]TGC-3’ motif, target of the Type I NgoAV RMS, in each of the 25 *N. gonorrhoeae* strains included in the study. Per-base distribution of IPD ratio is shown as two separate boxplots containing the values in the forward (red) and reverse (blue) strand.

Supplementary Figure 4. IPD ratio values for each base in all instances of the 5’-GCAN[8]TGC-3’ motif, target of the Type I NgoAV RMS, in each of the 25 *N. gonorrhoeae* strains included in the study. Per-base distribution of IPD ratio is shown as two separate boxplots containing the values in the forward (red) and reverse (blue) strand.

Supplementary Figure 5. IPD ratio values for each base in all instances of the 5’-GAGN[5]TAC-3’ motif, target of the Type I NgoAXVII RMS, in each of the 25 *N. gonorrhoeae* strains included in the study. Per-base distribution of IPD ratio is shown as two separate boxplots containing the values in the forward (red) and reverse (blue) strand.

Supplementary Figure 6. IPD ratios for methylated cytosines (mCs) in Type II RMS. The distribution of IPD ratios for the forward (red) and reverse (green) mCs as stated in REBASE are shown as boxplots together with a random 10k subsampling of unmethylated cytosines for each strain (blue). Statistical significance of the comparison among the distribution of the forward and reverse methylations with that from a random 10k subsampling of unmethylated cytosines is indicated next to the boxplots. ****p-value < 0.0001; ***p-value<0.001; **p-value<0.01;*p-value<0.05; ns: non-significant.

Supplementary Figure 7. IPD ratio values for each base in all instances of the 5’-AAANCGGTTNNC-3’ motif in each of the 25 *N. gonorrhoeae* strains included in the study. Per-base distribution of IPD ratio is shown as two separate boxplots containing the values in the forward (red) and reverse (blue) strand.

Supplementary Figure 8. IPD ratios for the underlined forward and reverse cytosines in the 5’-AAANCGGTTNNC-3’ motif. These values were compared to the distribution of a random 10k subsampling of unmethylated cytosines for each strain. Statistical significance is shown above the boxplots. ****p-value < 0.0001; ***p-value<0.001; **p-value<0.01;*p-value<0.05; ns: non-significant.

Supplementary Figure 9. IPD ratio values for each base in all instances of the 5’-GATC-3’ motif, target of the Type II NgoAXI RMS, in each of the 25 *N. gonorrhoeae* strains included in the study. Per-base distribution of IPD ratio is shown as two separate boxplots containing the values in the forward (red) and reverse (blue) strand.

Supplementary Figure 10. IPD ratio values for each base in all instances of the 5’-GGTGA-3’ motif, target of the Type II NgoAXVI RMS, in each of the 25 *N. gonorrhoeae* strains included in the study. Per-base distribution of IPD ratio is shown as two separate boxplots containing the values in the forward (red) and reverse (blue) strand.

Supplementary Figure 11. Inference of the methylated cytosine in the reverse strand of 5’-GGTGA-3’ (NgoAXVI). Bonferroni-corrected p-values are shown in logarithmic scale for the comparison between the distribution of IPD values for the underlined cytosine and a random 10k subsampling of unmethylated cytosines for each strain. The cytosine with a higher significance and thus, candidate to be methylated, is the second one.

Supplementary Figure 12. IPD ratio values for each base in all instances of the 5’-CCACC-3’ motif, target of the Type III NgoAX RMS, in each of the 25 *N. gonorrhoeae* strains included in the study. Per-base distribution of IPD ratio is shown as two separate boxplots containing the values in the forward (red) and reverse (blue) strand.

Supplementary Figure 13. IPD ratio values for each base in all instances of the 5’-GCAGA-3’ motif, target of the Type III NgoAXII RMS, in each of the 25 *N. gonorrhoeae* strains included in the study. Per-base distribution of IPD ratio is shown as two separate boxplots containing the values in the forward (red) and reverse (blue) strand.

Supplementary Figure 14. Maximum likelihood phylogenetic reconstruction of representative Dam protein sequences available in genomes of the *Neisseria* genus in the RefSeq database (accessed on 21st January 2019). These representative sequences (WP_*) were obtained using the Identical Protein Groups (IPG) tool in NCBI. Coloured strips represent the species in which these sequences are found. Those appearing only once have been coloured in grey as ‘Others’ but can be accessed using the ID on the tip labels in https://www.ncbi.nlm.nih.gov/ipg.

Supplementary Figure 15. Circular plot (58) representing the annotation of the WHO G pConjugative plasmid, which shows the presence of an orphan DNA methylase that has a 100% protein identity with two entries in REBASE.

## Acknowledgements

This work was supported by Wellcome grant number 098051 (LSB, SRH and JP) and the Foundation for Medical Research at Örebro University Hospital, Örebro, Sweden (DG and MU). We thank them for the financial support.

## Authors contributions statement

All co-authors planned and designed the study. LSB and SRH performed the bioinformatic analyses, with support from DG, MU and JP. LSB and SRH drafted the manuscript and all co-authors commented and approved the final version.

## Additional information

**Supplementary information** is available at XXX.

### Accession codes

Stated in the Data Availability section.

