## Supplementary Tables 1-2, 4-9 and Supplementary Figures 1-15 for "Genetic variation regulates the activation and specificity of Restriction-Modification systems in *Neisseria gonorrhoeae*"

This document contains Supplementary Tables 1-2, 4-9 and Supplementary Figures 1-15.

Supplementary Table 1. List of *N. gonorrhoeae* strains used in the analysis. The ENA accession information of the raw and complete genomic data are indicated along with the average per-base sequencing coverage. Sample accession numbers are used for raw data as some of the strains include more than one sequencing run.

| **Strain** | **PacBio raw data (ENA sample accession)** | **Genome data accession** | **Average per-base coverage (X)** | **References** |
|---|---|---|---|---|
| WHO F | SAMEA2448460 | LT591897 | 118.8 | (27) |
| WHO G | SAMEA2448461 | LT591898<br>LT591899<br>LT591900 | 120.4 | (27) |
| WHO K | SAMEA2448462 | LT591908<br>LT591909 | 173.0 | (27) |
| WHO L | SAMEA2448463 | LT591901<br>LT591902<br>LT591903 | 150.6 | (27) |
| WHO M | SAMEA2448464 | LT591904<br>LT591905<br>LT591906<br>LT591907 | 188.2 | (27) |
| WHO N | SAMEA2448465 | LT591910<br>LT591911<br>LT591912<br>LT591913 | 155.5 | (27) |
| WHO O | SAMEA2448466 | LT592146<br>LT592147<br>LT592148<br>LT592149 | 190.1 | (27) |
| WHO P | SAMEA2448467 | LT592157<br>LT592158 | 211.8 | (27) |
| WHO U | SAMEA2796327 | LT592159<br>LT592160 | 169.9 | (27) |
| WHO V | SAMEA2796328 | LT592150<br>LT592151<br>LT592152 | 142.2 | (27) |

| | | | | |
|---|---|---|---|---|
| WHO W | SAMEA2448470 | LT592163<br>LT592164<br>LT592165 | 132.5 | (27) |
| WHO X | SAMEA2448468 | LT592155<br>LT592156 | 204.6 | (27) |
| WHO Y | SAMEA2448469 | LT592161<br>LT592162 | 190.1 | (27) |
| WHO Z | SAMEA2796326 | LT592153<br>LT592154 | 163.7 | (27) |
| NCTC10928 | SAMEA3174297 | https://github.com/leosanbu/MethylationProject | 385.0 | NCTC 3000 |
| NCTC10931 | SAMEA3174298 | https://github.com/leosanbu/MethylationProject | 394.0 | NCTC 3000 |
| NCTC12700 | SAMEA3174299 | https://github.com/leosanbu/MethylationProject | 425.0 | NCTC 3000 |
| NCTC13795 | SAMEA4076737 | https://github.com/leosanbu/MethylationProject | 201.0 | NCTC 3000 |
| NCTC13798 | SAMEA4076741 | https://github.com/leosanbu/MethylationProject | 179.0 | NCTC 3000 |
| NCTC13799 | SAMEA4076765 | https://github.com/leosanbu/MethylationProject | 149.0 | NCTC 3000 |
| NCTC13800 | SAMEA4076768 | https://github.com/leosanbu/MethylationProject | 241.0 | NCTC 3000 |
| NCTC13801 | SAMEA4076769 | https://github.com/leosanbu/MethylationProject | 269.0 | NCTC 3000 |
| NCTC13802 | SAMEA4076770 | https://github.com/leosanbu/MethylationProject | 185.0 | NCTC 3000 |
| NCTC13805 | SAMEA4076773 | https://github.com/leosanbu/MethylationProject | 233.0 | NCTC 3000 |
| FA1090 | SAMN03775647 | NCBI:AE004969 | 367.0 | NCBI, (9) |

Supplementary Table 2. Results from a Gene Ontology (GO) enrichment analysis of the flanking genes of the 15 restriction-modifications system (RMS) in *N. gonorrhoeae*. The number of genes annotated in each GO term and the expected number of genes to fall in each category are shown together with the number of significant hits in the flanking genes. GO terms significantly enriched by both the *classic* and *weight01* algorithms (p-value < 0.05) are shown. Results are shown for the three sub-ontologies (BP = Biological Process, MF = Molecular Function, CC = Cellular Component).

| Ontology | GO term | Term | Annotated | Significant | Expected | classic (p-value) | weight01 (p-value) |
|---|---|---|---|---|---|---|---|
| BP | GO: 0006418 | tRNA aminoacylation for protein translation | 22 | 3 | 0.35 | 0.0045 | 0.0045 |
| BP | GO: 0006313 | transposition, DNA-mediated | 11 | 2 | 0.18 | 0.0123 | 0.0123 |
| MF | GO: 0004812 | aminoacyl-tRNA ligase activity | 22 | 3 | 0.29 | 0.0024 | 0.0024 |
| MF | GO: 0005524 | ATP binding | 170 | 6 | 2.21 | 0.0167 | 0.0167 |
| MF | GO: 0000287 | magnesium ion binding | 20 | 2 | 0.26 | 0.0264 | 0.0264 |
| MF | GO: 0016853 | isomerase activity | 57 | 4 | 0.74 | 0.0052 | 0.0417 |
| CC | GO: 0005737 | cytoplasm | 165 | 4 | 1.39 | 0.032 | 0.027 |
| CC | GO: 1902494 | catalytic complex | 35 | 2 | 0.3 | 0.032 | 0.032 |

Supplementary Table 4. Type I NgoAV restriction-modifications system (RMS). Active (A) or disrupted (D) components are indicated. Red highlight is used to mark a cause of disruption. The different sources of variability in the pattern recognition domain of the specificity unit, which cause different methylated motifs (in different colours) are shown. An active methyltransferase requires both the methylase and the specificity unit to be active. In that case, if the restriction enzyme is not functional, the methylase is tagged as 'orphan'. *hsdR*: Restriction endonuclease; *hsdM*: Methyltransferase; *hsdS*: Specificity unit.

| | **Components** | | | | | | | | |
|---|---|---|---|---|---|---|---|---|---|
| | | | | **Source of variability** | | | | | |
| **Strain** | ***hsdR*** | ***hsdM*** | ***hsdS*** | **Premature stop codon** | **Number of ATLE repeats** | **Middle frameshift** | **Orphan *hsdM*?** | **Active *hsdM*?** | **Methylated motif** |
| FA1090 | A | A | A | No | 2 | Yes | No | Yes | GCAN{8}TGC/GCAN{8}TGC |
| WHO F | D | A | D | No | 0 | Yes | No | No | - |
| WHO G | A | A | A | No | 2 | No | No | Yes | GACN{7}TGC/GCAN{7}GTC |
| WHO K | A | A | D | Yes | 2 | Yes | No | No | - |
| WHO L | A | D | D | Yes | 1 | Yes | No | No | - |
| WHO M | A | A | A | No | 1 | No | No | Yes | GACN{6}TGC/GCAN{6}GTC |
| WHO N | A | A | A | No | 2 | No | No | Yes | GACN{7}TGC/GCAN{7}GTC |
| WHO O | A | A | D | No | 0 | Yes | No | No | - |
| WHO P | A | D | D | Yes | 3 | Yes | No | No | - |
| WHO U | A | A | A | No | 2 | No | No | Yes | GACN{7}TGC/GCAN{7}GTC |
| WHO V | A | A | D | Yes | 1 | Yes | No | No | - |
| WHO W | A | A | A | No | 2 | Yes | No | Yes | GCAN{8}TGC/GCAN{8}TGC |
| WHO X | A | A | D | Yes | 2 | Yes | No | No | - |
| WHO Y | A | A | D | Yes | 2 | Yes | No | No | - |
| WHO Z | A | A | D | Yes | 2 | Yes | No | No | - |
| NCTC10928 | A | A | D | Yes | 1 | Yes | No | No | - |
| NCTC10931 | A | A | A | No | 1 | No | No | Yes | GACN{6}TGC/GCAN{6}GTC |
| NCTC12700 | A | A | A | No | 1 | No | No | Yes | Not detected |
| NCTC13795 | A | D | D | No | 1 | No | No | No | - |
| NCTC13798 | A | A | D | Yes | 2 | Yes | No | No | - |
| NCTC13799 | A | D | D | Yes | 2 | Yes | No | No | - |
| NCTC13800 | A | A | D | Yes | 1 | Yes | No | No | - |
| NCTC13801 | A | A | A | No | 2 | No | No | Yes | GACN{7}TGC/GCAN{7}GTC |
| NCTC13802 | A | A | D | Yes | 1 | Yes | No | No | - |
| NCTC13805 | A | A | D | Yes | 1 | Yes | No | No | - |

Supplementary Table 5. Type I NgoAXVIIP restriction-modifications system (RMS). Active (A) or disrupted (D, in red) components are indicated. An active methyltransferase requires both the methylase and the specificity unit to be active. In that case, if the restriction enzyme is not functional, the methylase is tagged as 'orphan' (blue). *hsdR*: Restriction endonuclease; *hsdM*: Methyltransferase; *hsdS*: Specificity unit.

| | Components | | | | | |
|---|---|---|---|---|---|---|
| **Strain** | ***hsdR*** | ***hsdM*** | ***hsdS*** | **Orphan *hsdM*?** | **Active *hsdM*?** | **Methylated motif** |
| FA1090 | A | A | A | No | Yes | GAGN{5}TAC/GTAN{5}CTC |
| WHO F | D | D | A | No | No | - |
| WHO G | A | A | A | No | Yes | GAGN{5}TAC/GTAN{5}CTC |
| WHO K | A | A | A | No | Yes | GAGN{5}TAC/GTAN{5}CTC |
| WHO L | D | A | A | Yes | Yes | GAGN{5}TAC/GTAN{5}CTC |
| WHO M | A | A | A | No | Yes | GAGN{5}TAC/GTAN{5}CTC |
| WHO N | A | A | A | No | Yes | GAGN{5}TAC/GTAN{5}CTC |
| WHO O | A | A | A | No | Yes | GAGN{5}TAC/GTAN{5}CTC |
| WHO P | A | A | A | No | Yes | GAGN{5}TAC/GTAN{5}CTC |
| WHO U | A | A | A | No | Yes | GAGN{5}TAC/GTAN{5}CTC |
| WHO V | A | A | A | No | Yes | GAGN{5}TAC/GTAN{5}CTC |
| WHO W | A | A | A | No | Yes | GAGN{5}TAC/GTAN{5}CTC |
| WHO X | A | A | A | No | Yes | GAGN{5}TAC/GTAN{5}CTC |
| WHO Y | A | A | A | No | Yes | GAGN{5}TAC/GTAN{5}CTC |
| WHO Z | A | A | A | No | Yes | GAGN{5}TAC/GTAN{5}CTC |
| NCTC10928 | D | A | A | Yes | Yes | GAGN{5}TAC/GTAN{5}CTC |
| NCTC10931 | D | A | A | Yes | Yes | GAGN{5}TAC/GTAN{5}CTC |
| NCTC12700 | A | A | A | No | Yes | GAGN{5}TAC/GTAN{5}CTC |
| NCTC13795 | D | A | A | Yes | Yes | GAGN{5}TAC/GTAN{5}CTC |
| NCTC13798 | A | A | A | No | Yes | GAGN{5}TAC/GTAN{5}CTC |
| NCTC13799 | A | A | A | No | Yes | GAGN{5}TAC/GTAN{5}CTC |
| NCTC13800 | A | A | A | No | Yes | GAGN{5}TAC/GTAN{5}CTC |
| NCTC13801 | D | A | A | Yes | Yes | GAGN{5}TAC/GTAN{5}CTC |
| NCTC13802 | D | A | D | No | No | - |
| NCTC13805 | D | A | D | No | No | - |

Supplementary Table 6. Type II restriction-modifications systems (RMSs). Active (A) or disrupted (D, in red) components are indicated. An active methyltransferase requires both the methylase and the specificity unit to be active. In that case, if the restriction enzyme is not functional, the methylase is tagged as 'orphan' (blue). R: Restriction enzyme; M: Methyltransferase. The Bonferroni-corrected p-values of a Mann-Whitney test between the IPD values of methylated cytosines of each motif (as indicated in REBASE) and the distribution of unmethylated cytosines for each strain are shown. ****p-value < 0.001; ***p-value<0.001; **p-value<0.01;*p-value<0.05; ns: non-significant.

| | | Components | | | | | Bonferroni-corrected p-values (Cytosine methylation) | | |
|---|---|---|---|---|---|---|---|---|---|
| **RM** | **Strain** | **R** | **M** | **Orphan M?** | **Active M?** | **Motif** | 5'-RGCGCY-3' | 3'-YCGCGR-5' | (5mC) |
| | FA1090 | A | A | No | Yes | RGCGCY | 9.47E-93 | 1.03E-93 | **** |
| | WHO F | A | A | No | Yes | RGCGCY | 2.17E-96 | 1.42E-101 | **** |
| | WHO G | A | A | No | Yes | RGCGCY | 1.75E-80 | 2.70E-85 | **** |
| | WHO K | A | A | No | Yes | RGCGCY | 1.94E-95 | 2.62E-114 | **** |
| | WHO L | A | A | No | Yes | RGCGCY | 6.70E-86 | 1.24E-95 | **** |
| | WHO M | A | A | No | Yes | RGCGCY | 3.40E-110 | 1.86E-103 | **** |
| | WHO N | A | A | No | Yes | RGCGCY | 1.07E-97 | 2.24E-93 | **** |
| | WHO O | A | A | No | Yes | RGCGCY | 5.19E-114 | 2.95E-108 | **** |
| | WHO P | A | A | No | Yes | RGCGCY | 1.88E-107 | 1.94E-113 | **** |
| | WHO U | A | A | No | Yes | RGCGCY | 1.05E-87 | 6.28E-101 | **** |
| | WHO V | A | A | No | Yes | RGCGCY | 1.44E-98 | 4.08E-98 | **** |
| | WHO W | A | A | No | Yes | RGCGCY | 5.43E-93 | 1.54E-99 | **** |
| NgoAI | WHO X | A | A | No | Yes | RGCGCY | 2.01E-103 | 8.49E-97 | **** |
| | WHO Y | A | A | No | Yes | RGCGCY | 2.51E-116 | 5.78E-108 | **** |
| | WHO Z | A | A | No | Yes | RGCGCY | 1.52E-102 | 2.98E-97 | **** |
| | NCTC10928 | A | A | No | Yes | RGCGCY | 1.56E-116 | 7.33E-121 | **** |
| | NCTC10931 | A | A | No | Yes | RGCGCY | 1.56E-123 | 2.17E-123 | **** |
| | NCTC12700 | A | A | No | Yes | RGCGCY | 5.30E-127 | 8.16E-114 | **** |
| | NCTC13795 | A | A | No | Yes | RGCGCY | 2.32E-125 | 1.95E-121 | **** |
| | NCTC13798 | A | A | No | Yes | RGCGCY | 6.24E-130 | 3.65E-129 | **** |
| | NCTC13799 | A | A | No | Yes | RGCGCY | 1.48E-109 | 2.78E-121 | **** |
| | NCTC13800 | A | A | No | Yes | RGCGCY | 2.40E-130 | 2.60E-130 | **** |
| | NCTC13801 | A | A | No | Yes | RGCGCY | 1.81E-131 | 4.29E-124 | **** |
| | NCTC13802 | A | A | No | Yes | RGCGCY | 1.31E-120 | 1.83E-136 | **** |
| | NCTC13805 | A | A | No | Yes | RGCGCY | 9.18E-113 | 3.27E-103 | **** |
| **RM** | **Strain** | **R** | **M** | **Orphan M?** | **Active M?** | **Motif** | 5'-GGCC-3' | 3'-CCGG-5' | (5mC) |
| | FA1090 | D | A | Yes | Yes | GGCC | 6.84E-124 | 1.59E-120 | **** |
| | WHO F | D | A | Yes | Yes | GGCC | 7.19E-189 | 3.33E-194 | **** |
| | WHO G | A | A | No | Yes | GGCC | 3.60E-297 | 1.42E-305 | **** |
| | WHO K | D | A | Yes | Yes | GGCC | 5.44E-224 | 2.08E-238 | **** |
| | WHO L | D | A | Yes | Yes | GGCC | 3.29E-242 | 1.42E-248 | **** |
| | WHO M | D | A | Yes | Yes | GGCC | 1.08E-249 | 6.61E-286 | **** |
| | WHO N | A | A | No | Yes | GGCC | 2.12E-235 | 7.17E-265 | **** |
| | WHO O | A | A | No | Yes | GGCC | 2.79E-261 | 3.24E-245 | **** |
| | WHO P | D | A | Yes | Yes | GGCC | 1.12E-298 | 0.00E+00 | **** |
| | WHO U | D | A | Yes | Yes | GGCC | 7.19E-246 | 6.03E-252 | **** |
| | WHO V | D | A | Yes | Yes | GGCC | 5.13E-246 | 8.20E-270 | **** |
| | WHO W | D | A | Yes | Yes | GGCC | 8.24E-256 | 4.83E-281 | **** |
| NgoAII | WHO X | D | A | Yes | Yes | GGCC | 1.60E-280 | 6.43E-305 | **** |
| | WHO Y | D | A | Yes | Yes | GGCC | 0.00E+00 | 0.00E+00 | **** |
| | WHO Z | D | A | Yes | Yes | GGCC | 8.33E-229 | 2.52E-243 | **** |
| | NCTC10928 | A | A | No | Yes | GGCC | 2.17E-73 | 2.09E-70 | **** |
| | NCTC10931 | D | A | Yes | Yes | GGCC | 8.10E-82 | 9.35E-75 | **** |
| | NCTC12700 | A | A | No | Yes | GGCC | 5.05E-90 | 2.10E-76 | **** |
| | NCTC13795 | A | A | No | Yes | GGCC | 3.15E-105 | 2.62E-80 | **** |
| | NCTC13798 | D | A | Yes | Yes | GGCC | 8.32E-112 | 7.41E-95 | **** |
| | NCTC13799 | D | A | Yes | Yes | GGCC | 3.70E-125 | 1.24E-126 | **** |
| | NCTC13800 | D | A | Yes | Yes | GGCC | 1.75E-119 | 2.18E-114 | **** |
| | NCTC13801 | A | A | No | Yes | GGCC | 3.23E-121 | 1.68E-102 | **** |
| | NCTC13802 | D | A | Yes | Yes | GGCC | 1.07E-93 | 4.38E-80 | **** |

| | | | | | | | | | |
|---|---|---|---|---|---|---|---|---|---|
| | NCTC13805 | D | A | Yes | Yes | GGCC | 3.50E-71 | 5.14E-91 | **** |
| **RM** | **Strain** | **R** | **M** | **Orphan M?** | **Active M?** | **Motif** | 5'-CCGCGG-3' | 3'-GGCGCC-5' | (5mC) |
| | FA1090 | A | A | No | Yes | CCGCGG | 1.30E-23 | 2.99E-19 | **** |
| | WHO F | Absent | | No | No | - | 1 | 1 | ns |
| | WHO G | A | A | No | Yes | CCGCGG | 6.03E-32 | 1.36E-33 | **** |
| | WHO K | A | A | No | Yes | CCGCGG | 3.72E-45 | 4.85E-36 | **** |
| | WHO L | Absent | | No | No | - | 0.854309627 | 1 | ns |
| | WHO M | A | A | No | Yes | CCGCGG | 5.92E-33 | 7.52E-34 | **** |
| | WHO N | A | A | No | Yes | CCGCGG | 8.05E-34 | 8.89E-25 | **** |
| | WHO O | A | A | No | Yes | CCGCGG | 7.29E-34 | 1.23E-25 | **** |
| | WHO P | A | A | No | Yes | CCGCGG | 9.71E-39 | 9.15E-36 | **** |
| | WHO U | A | A | No | Yes | CCGCGG | 8.22E-23 | 6.74E-26 | **** |
| | WHO V | A | A | No | Yes | CCGCGG | 2.11E-30 | 1.97E-29 | **** |
| | WHO W | A | A | No | Yes | CCGCGG | 3.34E-34 | 4.23E-33 | **** |
| NgoAIII | WHO X | A | A | No | Yes | CCGCGG | 4.70E-43 | 2.94E-38 | **** |
| | WHO Y | A | A | No | Yes | CCGCGG | 2.84E-43 | 1.21E-37 | **** |
| | WHO Z | A | A | No | Yes | CCGCGG | 5.06E-28 | 7.33E-19 | **** |
| | NCTC10928 | Absent | | No | No | - | 1 | 0.125551747 | ns |
| | NCTC10931 | A | A | No | Yes | CCGCGG | 2.20E-01 | 0.031189686 | ns/* |
| | NCTC12700 | Absent | | No | No | - | 2.29E-06 | 0.000732929 | ****/*** |
| | NCTC13795 | A | A | No | Yes | CCGCGG | 2.21E-02 | 0.001653781 | */** |
| | NCTC13798 | A | A | No | Yes | CCGCGG | 1.81E-03 | 0.023353729 | **/* |
| | NCTC13799 | A | A | No | Yes | CCGCGG | 1.12E-02 | 0.003917145 | */** |
| | NCTC13800 | A | A | No | Yes | CCGCGG | 1.05E-03 | 0.000816589 | **/*** |
| | NCTC13801 | A | A | No | Yes | CCGCGG | 5.84E-04 | 0.000900113 | *** |
| | NCTC13802 | A | A | No | Yes | CCGCGG | 4.41E-04 | 0.002522283 | ***/** |
| | NCTC13805 | A | A | No | Yes | CCGCGG | 1.69E-01 | 0.024751118 | ns/* |
| **RM** | **Strain** | **R** | **M** | **Orphan M?** | **Active M?** | **Motif** | 5'-GCCGGC-3' | 3'-CGGCCG-5' | (5mC) |
| | FA1090 | A | A | No | Yes | GCCGGC | 1.77E-271 | 4.52E-263 | **** |
| | WHO F | A | A | No | Yes | GCCGGC | 3.53E-203 | 3.75E-218 | **** |
| | WHO G | A | A | No | Yes | GCCGGC | 1.26E-289 | 1.79E-272 | **** |
| | WHO K | A | A | No | Yes | GCCGGC | 2.23E-253 | 1.30E-229 | **** |
| | WHO L | A | A | No | Yes | GCCGGC | 5.89E-226 | 2.81E-270 | **** |
| | WHO M | A | A | No | Yes | GCCGGC | 7.95E-274 | 3.23E-271 | **** |
| | WHO N | A | A | No | Yes | GCCGGC | 1.19E-261 | 8.17E-253 | **** |
| | WHO O | A | A | No | Yes | GCCGGC | 2.99E-280 | 2.44E-265 | **** |
| | WHO P | A | A | No | Yes | GCCGGC | 0.00E+00 | 0.00E+00 | **** |
| | WHO U | A | A | No | Yes | GCCGGC | 0.00E+00 | 0.00E+00 | **** |
| | WHO V | A | A | No | Yes | GCCGGC | 1.67E-268 | 2.02E-268 | **** |
| | WHO W | A | A | No | Yes | GCCGGC | 2.24E-276 | 4.76E-246 | **** |
| NgoAIV | WHO X | A | A | No | Yes | GCCGGC | 2.00E-290 | 1.13E-284 | **** |
| | WHO Y | A | A | No | Yes | GCCGGC | 0.00E+00 | 0.00E+00 | **** |
| | WHO Z | A | A | No | Yes | GCCGGC | 0.00E+00 | 0.00E+00 | **** |
| | NCTC10928 | A | A | No | Yes | GCCGGC | 0.00E+00 | 0.00E+00 | **** |
| | NCTC10931 | A | A | No | Yes | GCCGGC | 0.00E+00 | 0.00E+00 | **** |
| | NCTC12700 | A | A | No | Yes | GCCGGC | 0.00E+00 | 0.00E+00 | **** |
| | NCTC13795 | A | A | No | Yes | GCCGGC | 0.00E+00 | 0.00E+00 | **** |
| | NCTC13798 | A | A | No | Yes | GCCGGC | 0.00E+00 | 0.00E+00 | **** |
| | NCTC13799 | A | A | No | Yes | GCCGGC | 0.00E+00 | 0.00E+00 | **** |
| | NCTC13800 | A | A | No | Yes | GCCGGC | 0.00E+00 | 0.00E+00 | **** |
| | NCTC13801 | A | A | No | Yes | GCCGGC | 0.00E+00 | 0.00E+00 | **** |
| | NCTC13802 | A | A | No | Yes | GCCGGC | 0.00E+00 | 0.00E+00 | **** |
| | NCTC13805 | A | A | No | Yes | GCCGGC | 0.00E+00 | 0.00E+00 | **** |

| RM | Strain | R | M | Orphan M? | Active M? | Motif | 5'-GCGGC-3' | 3'-CGCCG-5' | (5mC) |
|---|---|---|---|---|---|---|---|---|---|
| | FA1090 | A | A | No | Yes | GCGGC | 7.56E-123 | 6.17E-68 | **** |
| | WHO F | A | A | No | Yes | GCGGC | 3.78E-92 | 3.32E-22 | **** |
| | WHO G | A | A | No | Yes | GCGGC | 1.00E+00 | 3.72E-58 | ns/**** |
| | WHO K | A | A | No | Yes | GCGGC | 7.09E-119 | 5.88E-48 | **** |
| | WHO L | A | A | No | Yes | GCGGC | 1.06E-54 | 9.90E-63 | **** |
| | WHO M | A | A | No | Yes | GCGGC | 5.88E-54 | 2.33E-48 | **** |
| | WHO N | A | A | No | Yes | GCGGC | 2.46E-49 | 2.83E-51 | **** |
| | WHO O | A | A | No | Yes | GCGGC | 6.76E-120 | 1.67E-65 | **** |
| | WHO P | A | A | No | Yes | GCGGC | 7.85E-102 | 2.67E-74 | **** |
| | WHO U | A | A | No | Yes | GCGGC | 4.22E-19 | 6.56E-108 | **** |
| | WHO V | A | A | No | Yes | GCGGC | 1.87E-85 | 5.11E-103 | **** |
| | WHO W | A | A | No | Yes | GCGGC | 8.73E-148 | 3.31E-89 | **** |
| NgoAVII | WHO X | A | A | No | Yes | GCGGC | 1.90E-95 | 6.45E-42 | **** |
| | WHO Y | A | A | No | Yes | GCGGC | 0.00E+00 | 0.00E+00 | **** |
| | WHO Z | A | A | No | Yes | GCGGC | 0.00E+00 | 0.00E+00 | **** |
| | NCTC10928 | A | A | No | Yes | GCGGC | 2.40E-234 | 0.00E+00 | **** |
| | NCTC10931 | A | A | No | Yes | GCGGC | 5.16E-237 | 0.00E+00 | **** |
| | NCTC12700 | A | A | No | Yes | GCGGC | 0.00E+00 | 0.00E+00 | **** |
| | NCTC13795 | A | A | No | Yes | GCGGC | 2.07E-289 | 0.00E+00 | **** |
| | NCTC13798 | A | A | No | Yes | GCGGC | 0.00E+00 | 0.00E+00 | **** |
| | NCTC13799 | A | A | No | Yes | GCGGC | 0.00E+00 | 0.00E+00 | **** |
| | NCTC13800 | A | A | No | Yes | GCGGC | 0.00E+00 | 0.00E+00 | **** |
| | NCTC13801 | A | A | No | Yes | GCGGC | 0.00E+00 | 0.00E+00 | **** |
| | NCTC13802 | A | A | No | Yes | GCGGC | 0.00E+00 | 0.00E+00 | **** |
| | NCTC13805 | A | A | No | Yes | GCGGC | 0.00E+00 | 0.00E+00 | **** |

| RM | Strain | RM | S | Orphan M? | Active M? | Motif |
|---|---|---|---|---|---|---|
| | FA1090 | A | A | No | Yes | Unknown |
| | WHO F | A | A | No | Yes | Unknown |
| | WHO G | A | A | No | Yes | Unknown |
| | WHO K | A | A | No | Yes | Unknown |
| | WHO L | A | A | No | Yes | Unknown |
| | WHO M | A | A | No | Yes | Unknown |
| | WHO N | A | A | No | Yes | Unknown |
| | WHO O | A | A | No | Yes | Unknown |
| | WHO P | A | A | No | Yes | Unknown |
| | WHO U | A | A | No | Yes | Unknown |
| | WHO V | A | A | No | Yes | Unknown |
| | WHO W | A | A | No | Yes | Unknown |
| NgoAVIII | WHO X | A | A | No | Yes | Unknown |
| | WHO Y | A | A | No | Yes | Unknown |
| | WHO Z | A | A | No | Yes | Unknown |
| | NCTC10928 | A | A | No | Yes | Unknown |
| | NCTC10931 | A | A | No | Yes | Unknown |
| | NCTC12700 | A | A | No | Yes | Unknown |
| | NCTC13795 | A | A | No | Yes | Unknown |
| | NCTC13798 | A | A | No | Yes | Unknown |
| | NCTC13799 | A | A | No | Yes | Unknown |
| | NCTC13800 | A | A | No | Yes | Unknown |
| | NCTC13801 | A | A | No | Yes | Unknown |
| | NCTC13802 | A | A | No | Yes | Unknown |
| | NCTC13805 | A | A | No | Yes | Unknown |

| RM | Strain | VSR | M | Orphan M? | Active M? | Motif |
|---|---|---|---|---|---|---|
| NgoAXIII | FA1090 | - | D | No | No | - |
| | WHO F | A | D | No | No | - |
| | WHO G | A | D | No | No | - |
| | WHO K | A | D | No | No | - |
| | WHO L | D | D | No | No | - |
| | WHO M | A | D | No | No | - |
| | WHO N | A | D | No | No | - |
| | WHO O | A | D | No | No | - |
| | WHO P | A | D | No | No | - |
| | WHO U | A | D | No | No | - |
| | WHO V | A | D | No | No | - |
| | WHO W | A | D | No | No | - |
| | WHO X | A | D | No | No | - |
| | WHO Y | A | D | No | No | - |
| | WHO Z | A | D | No | No | - |
| | NCTC10928 | A | D | No | No | - |
| | NCTC10931 | A | D | No | No | - |
| | NCTC12700 | A | D | No | No | - |
| | NCTC13795 | A | D | No | No | - |
| | NCTC13798 | A | D | No | No | - |
| | NCTC13799 | A | D | No | No | - |
| | NCTC13800 | A | D | No | No | - |
| | NCTC13801 | A | D | No | No | - |
| | NCTC13802 | A | D | No | No | - |
| | NCTC13805 | A | D | No | No | - |

| RM | Strain | R | M | Orphan M? | Active M? | Motif |
|---|---|---|---|---|---|---|
| NgoAXIP | FA1090 | A | D | No | No | - |
| | WHO F | A | D | No | No | - |
| | WHO G | A | D | No | No | - |
| | WHO K | A | D | No | No | - |
| | WHO L | A | D | No | No | - |
| | WHO M | A | D | No | No | - |
| | WHO N | A | D | No | No | - |
| | WHO O | A | D | No | No | - |
| | WHO P | A | D | No | No | - |
| | WHO U | A | D | No | No | - |
| | WHO V | A | D | No | No | - |
| | WHO W | A | D | No | No | - |
| | WHO X | A | D | No | No | - |
| | WHO Y | A | D | No | No | - |
| | WHO Z | A | D | No | No | - |
| | NCTC10928 | A | D | No | No | - |
| | NCTC10931 | Dam | D | Yes | Yes | GATC |
| | NCTC12700 | Dam | D | Yes | Yes | GATC |
| | NCTC13795 | A | D | No | No | - |
| | NCTC13798 | A | D | No | No | - |
| | NCTC13799 | A | D | No | No | - |
| | NCTC13800 | A | D | No | No | - |
| | NCTC13801 | A | D | No | No | - |
| | NCTC13802 | A | D | No | No | - |
| | NCTC13805 | A | D | No | No | - |

| RM | Strain | VSR | M | Orphan M? | Active M? | Motif | 5'-CCGG-3' | 3-GGCC-5' | |
|---|---|---|---|---|---|---|---|---|---|
| | FA1090 | A | A | No | Yes | CCGG | 1.95E-41 | 6.38E-47 | **** |
| | WHO F | A | A | No | Yes | CCGG | 1.47E-120 | 1.71E-124 | **** |
| | WHO G | A | A | No | Yes | CCGG | 5.05E-29 | 2.68E-28 | **** |
| | WHO K | A | A | No | Yes | CCGG | 3.37E-119 | 1.03E-117 | **** |
| | WHO L | A | A | No | Yes | CCGG | 8.86E-56 | 1.53E-94 | **** |
| | WHO M | A | A | No | Yes | CCGG | 7.51E-79 | 1.15E-92 | **** |
| | WHO N | A | A | No | Yes | CCGG | 2.21E-51 | 4.35E-59 | **** |
| | WHO O | A | A | No | Yes | CCGG | 4.09E-123 | 1.30E-119 | **** |
| | WHO P | A | A | No | Yes | CCGG | 3.27E-107 | 2.54E-111 | **** |
| | WHO U | A | A | No | Yes | CCGG | 0.040205158 | 0.005377179 | */** |
| | WHO V | A | A | No | Yes | CCGG | 1.63E-19 | 4.60E-21 | **** |
| | WHO W | A | A | No | Yes | CCGG | 1.52E-64 | 3.32E-82 | **** |
| NgoAXIV | WHO X | A | A | No | Yes | CCGG | 3.61E-143 | 7.17E-151 | **** |
| | WHO Y | A | A | No | Yes | CCGG | 3.30E-129 | 1.47E-144 | **** |
| | WHO Z | A | A | No | Yes | CCGG | 3.78E-07 | 9.17E-14 | **** |
| | NCTC10928 | A | A | No | Yes | CCGG | 2.02E-36 | 2.12E-47 | **** |
| | NCTC10931 | A | A | No | Yes | CCGG | 1.38E-27 | 9.10E-45 | **** |
| | NCTC12700 | A | A | No | Yes | CCGG | 7.13E-44 | 3.96E-50 | **** |
| | NCTC13795 | A | A | No | Yes | CCGG | 1.05E-62 | 7.93E-75 | **** |
| | NCTC13798 | A | A | No | Yes | CCGG | 9.85E-54 | 1.69E-78 | **** |
| | NCTC13799 | A | A | No | Yes | CCGG | 2.30E-105 | 1.38E-123 | **** |
| | NCTC13800 | A | A | No | Yes | CCGG | 8.55E-73 | 6.82E-105 | **** |
| | NCTC13801 | A | A | No | Yes | CCGG | 5.43E-72 | 8.88E-79 | **** |
| | NCTC13802 | A | A | No | Yes | CCGG | 2.99E-89 | 1.35E-63 | **** |
| | NCTC13805 | A | A | No | Yes | CCGG | 8.87E-55 | 1.01E-81 | **** |

| RM | Strain | R | M | Orphan M? | Active M? | Motif | 5'-GGNNCC-3' | 3-CCNNGG-5' | (5mC) |
|---|---|---|---|---|---|---|---|---|---|
| | FA1090 | A | A | No | Yes | GGNNCC | 8.84E-252 | 1.46E-234 | **** |
| | WHO F | A | A | No | Yes | GGNNCC | 1.05E-200 | 5.33E-191 | **** |
| | WHO G | D | A | Yes | Yes | GGNNCC | 0.00E+00 | 0.00E+00 | **** |
| | WHO K | A | A | No | Yes | GGNNCC | 1.23E-226 | 6.68E-209 | **** |
| | WHO L | A | A | No | Yes | GGNNCC | 2.33E-207 | 3.27E-242 | **** |
| | WHO M | D | A | Yes | Yes | GGNNCC | 4.70E-264 | 2.45E-252 | **** |
| | WHO N | D | A | Yes | Yes | GGNNCC | 6.64E-223 | 1.54E-245 | **** |
| | WHO O | D | A | Yes | Yes | GGNNCC | 1.42E-247 | 4.20E-237 | **** |
| | WHO P | D | A | Yes | Yes | GGNNCC | 7.06E-291 | 1.11E-284 | **** |
| | WHO U | D | A | Yes | Yes | GGNNCC | 2.60E-283 | 6.29E-280 | **** |
| | WHO V | A | A | No | Yes | GGNNCC | 1.15E-259 | 8.75E-266 | **** |
| | WHO W | A | A | No | Yes | GGNNCC | 2.18E-234 | 2.95E-231 | **** |
| NgoAXV | WHO X | A | A | No | Yes | GGNNCC | 1.14E-240 | 2.58E-258 | **** |
| | WHO Y | A | A | No | Yes | GGNNCC | 1.51E-279 | 1.35E-297 | **** |
| | WHO Z | A | A | No | Yes | GGNNCC | 1.78E-245 | 3.09E-267 | **** |
| | NCTC10928 | A | A | No | Yes | GGNNCC | 7.72E-134 | 2.68E-158 | **** |
| | NCTC10931 | A | A | No | Yes | GGNNCC | 1.64E-152 | 4.17E-168 | **** |
| | NCTC12700 | A | A | No | Yes | GGNNCC | 1.27E-153 | 1.73E-166 | **** |
| | NCTC13795 | A | A | No | Yes | GGNNCC | 3.19E-154 | 1.77E-159 | **** |
| | NCTC13798 | A | A | No | Yes | GGNNCC | 6.07E-164 | 2.31E-168 | **** |
| | NCTC13799 | D | A | Yes | Yes | GGNNCC | 4.22E-170 | 5.46E-186 | **** |
| | NCTC13800 | A | A | No | Yes | GGNNCC | 4.09E-179 | 3.20E-175 | **** |
| | NCTC13801 | A | A | No | Yes | GGNNCC | 1.29E-168 | 2.33E-170 | **** |
| | NCTC13802 | A | A | No | Yes | GGNNCC | 1.45E-157 | 7.76E-148 | **** |
| | NCTC13805 | A | A | No | Yes | GGNNCC | 6.37E-135 | 3.72E-160 | **** |

| RM | Strain | R | M | Orphan M? | Active M? | Motif |
|---|---|---|---|---|---|---|
| NgoAXVI | FA1090 | D | A | Yes | Yes | GGTGA |
| | WHO F | D | A | Yes | Yes | GGTGA |
| | WHO G | A | A | No | Yes | GGTGA |
| | WHO K | A | A | No | Yes | GGTGA |
| | WHO L | A | A | No | Yes | GGTGA |
| | WHO M | A | A | No | Yes | GGTGA |
| | WHO N | A | A | No | Yes | GGTGA |
| | WHO O | D | A | Yes | Yes | GGTGA |
| | WHO P | A | A | No | Yes | GGTGA |
| | WHO U | A | A | No | Yes | GGTGA |
| | WHO V | A | A | No | Yes | GGTGA |
| | WHO W | A | A | No | Yes | GGTGA |
| | WHO X | A | A | No | Yes | GGTGA |
| | WHO Y | A | A | No | Yes | GGTGA |
| | WHO Z | A | A | No | Yes | GGTGA |
| | NCTC10928 | A | A | No | Yes | GGTGA |
| | NCTC10931 | A | A | No | Yes | GGTGA |
| | NCTC12700 | A | A | No | Yes | GGTGA |
| | NCTC13795 | A | A | No | Yes | GGTGA |
| | NCTC13798 | A | A | No | Yes | GGTGA |
| | NCTC13799 | A | A | No | Yes | GGTGA |
| | NCTC13800 | A | A | No | Yes | GGTGA |
| | NCTC13801 | A | A | No | Yes | GGTGA |
| | NCTC13802 | A | A | No | Yes | GGTGA |
| | NCTC13805 | A | A | No | Yes | GGTGA |

Supplementary Table 7. Type III NgoAX restriction-modifications system (RMS). Active (A) or disrupted (D, in red) components are indicated. The number of CCCAA repeats and the DNA recognition domain (DRD) allele are shown, which are the source of variability in the Mod unit. An active methyltransferase requires both the methylase and the specificity unit to be active. In that case, if the restriction enzyme is not functional, the methylase is tagged as 'orphan'. *Res* = Restriction enzyme, *Mod* = Methyltransferase.

| | | | Source of variability | | | | |
|---|---|---|---|---|---|---|---|
| **Strain** | ***Res*** | ***Mod*** | **Number of CCCAA repeats** | **DRD allele** | **Orphan *Mod*?** | **Active *Mod*?** | **Motif** |
| FA1090 | A | A | 12 | *modB1* | No | Yes | CCACC |
| WHO F | A | A | 6 | *modB1* | No | Yes | CCACC |
| WHO G | A | D | 13 | *modB1* | No | No | - |
| WHO K | A | D | 10 | *modB1* | No | No | - |
| WHO L | A | D | 5 | *modB1* | No | No | - |
| WHO M | A | D | 8 | *modB1* | No | No | - |
| WHO N | A | D | 13 | *modB1* | No | No | - |
| WHO O | A | D | 7 | *modB1* | No | No | - |
| WHO P | A | A | 18 | *modB1* | No | Yes | CCACC |
| WHO U | A | D | 8 | *modB1* | No | No | - |
| WHO V | A | D | 10 | *modB1* | No | No | - |
| WHO W | A | D | 8 | *modB1* | No | No | - |
| WHO X | A | D | 8 | *modB1* | No | No | - |
| WHO Y | A | A | 9 | *modB1* | No | Yes | CCACC |
| WHO Z | A | D | 5 | *modB1* | No | No | - |
| NCTC10928 | A | D | 19 | *modB1* | No | No | - |
| NCTC10931 | A | D | 7 | *modB1* | No | No | - |
| NCTC12700 | A | A | 36 | *modB1* | No | Yes | CCACC |
| NCTC13795 | A | D | 7 | *modB1* | No | No | - |
| NCTC13798 | A | A | 9 | *modB1* | No | Yes | CCACC |
| NCTC13799 | A | D | 4 | *modB1* | No | No | - |
| NCTC13800 | A | D | 7 | *modB1* | No | No | - |
| NCTC13801 | A | D | 10 | *modB1* | No | No | - |
| NCTC13802 | A | D | 7 | *modB1* | No | No | - |
| NCTC13805 | A | D | 7 | *modB1* | No | No | - |

Supplementary Table 8. Type III NgoAXII restriction-modifications system (RMS). Active (A) or disrupted (D, in red) components are indicated. The number of AGCC repeats and the DNA-recognition domain (DRD) allele are shown, which are the source of variability in the Mod unit. An active methyltransferase requires both the methylase and the specificity unit to be active. In that case, if the restriction enzyme is not functional, the methylase is tagged as 'orphan'. *Res* = Restriction enzyme, *Mod* = Methyltransferase.

| Strain | *Res* | *Mod* | Source of variability | | Orphan *Mod*? | Active *Mod*? | Motif |
|---|---|---|---|---|---|---|---|
| | | | Number of AGCC repeats | DRD allele | | | |
| FA1090 | A | D | 35 | *modA13* | No | No | - |
| WHO F | A | D | 22 | *modA13* | No | No | - |
| WHO G | A | D | 14 | *modA13* | No | No | - |
| WHO K | A | D | 23 | *modA13* | No | No | - |
| WHO L | A | D | 14 | *modA13* | No | No | - |
| WHO M | A | D | 12 | *modA13* | No | No | - |
| WHO N | A | A | 28 | *modA13* | No | Yes | GCAGA |
| WHO O | A | D | 15 | *modA13* | No | No | - |
| WHO P | A | D | 15 | *modA13* | No | No | - |
| WHO U | A | D | 15 | *modA13* | No | No | - |
| WHO V | A | D | 18 | *modA13* | No | No | - |
| WHO W | A | D | 20 | *modA13* | No | No | - |
| WHO X | A | D | 21 | *modA13* | No | No | - |
| WHO Y | A | D | 18 | *modA13* | No | No | - |
| WHO Z | A | D | 18 | *modA13* | No | No | - |
| NCTC10928 | A | D | 14 | *modA13* | No | No | - |
| NCTC10931 | A | D | 18 | *modA13* | No | No | - |
| NCTC12700 | A | D | 15 | *modA13* | No | No | - |
| NCTC13795 | A | D | 25 | *modA13* | No | No | - |
| NCTC13798 | A | A | 13 | *modA13* | No | Yes | GCAGA |
| NCTC13799 | A | A | 19 | *modA13* | No | Yes | GCAGA |
| NCTC13800 | A | D | 17 | *modA13* | No | No | - |
| NCTC13801 | A | D | 12 | *modA13* | No | No | - |
| NCTC13802 | A | D | 3 | *modA13* | No | No | - |
| NCTC13805 | A | D | 3 | *modA13* | No | No | - |

Supplementary Table 9. Number of GCAGA and AGAAA and GCAGAAATTY motifs in each strain and number and proportion of those with an IPD>3 in the following underlined bases. GCAGA is the motif associated to NgoAXII detected by the PacBio SMRT analysis pipeline (NCTC13798, NCTC13799 and WHO N), AGAAA is the predicted motif for this methylase in REBASE and GCAGAAATTY is the motif resulting from overlapping GCAGA, AGAAA and the recognition sequence of the ApoI enzyme (5'-RAATTY-3').

| | GCAG**A** | | | AGA**A**A | | | GCAGAAATTY | | | | |
|---|---|---|---|---|---|---|---|---|---|---|---|
| | Number of motifs | Number of motifs with IPD>3 in GCAGA | % methylated motifs (IPD>3) | Number of motifs | Number of motifs with IPD>3 in AGAAA | % methylated motifs (IPD>3) | Number of motifs | Number of motifs with IPD>3 in GCAG**A**AATTY | % methylated GCAG**A**AATTY (IPD>3) | Number of motifs with IPD>3 in GCAGA**A**ATTY | % methylated GCAGA**A**ATTY (IPD>3) |
| FA1090 | 4560 | 3 | 0.07 | 5134 | 0 | 0.00 | 5 | 0 | 0.00 | 0 | 0.00 |
| NCTC10928 | 4779 | 9 | 0.19 | 5449 | 1 | 0.02 | 5 | 0 | 0.00 | 0 | 0.00 |
| NCTC10931 | 4984 | 111 | 2.23 | 5938 | 0 | 0.00 | 7 | 0 | 0.00 | 0 | 0.00 |
| NCTC12700 | 4628 | 77 | 1.66 | 5250 | 0 | 0.00 | 5 | 0 | 0.00 | 0 | 0.00 |
| NCTC13795 | 4672 | 12 | 0.26 | 5337 | 2 | 0.04 | 6 | 0 | 0.00 | 0 | 0.00 |
| NCTC13798 | 4746 | 3970 | 83.65 | 5507 | 1 | 0.02 | 6 | 4 | 66.67 | 0 | 0.00 |
| NCTC13799 | 4611 | 4169 | 90.41 | 5288 | 2 | 0.04 | 5 | 3 | 60.00 | 0 | 0.00 |
| NCTC13800 | 4740 | 10 | 0.21 | 5498 | 0 | 0.00 | 5 | 0 | 0.00 | 0 | 0.00 |
| NCTC13801 | 4739 | 32 | 0.68 | 5483 | 0 | 0.00 | 5 | 0 | 0.00 | 0 | 0.00 |
| NCTC13802 | 4842 | 10 | 0.21 | 5568 | 0 | 0.00 | 8 | 0 | 0.00 | 0 | 0.00 |
| NCTC13805 | 5110 | 11 | 0.22 | 5938 | 1 | 0.02 | 7 | 0 | 0.00 | 0 | 0.00 |
| WHO F | 4856 | 2 | 0.04 | 5728 | 0 | 0.00 | 8 | 0 | 0.00 | 0 | 0.00 |
| WHO G | 4742 | 29 | 0.61 | 5447 | 0 | 0.00 | 4 | 0 | 0.00 | 0 | 0.00 |
| WHO K | 4615 | 8 | 0.17 | 5254 | 0 | 0.00 | 5 | 0 | 0.00 | 0 | 0.00 |
| WHO L | 4737 | 9 | 0.19 | 5360 | 0 | 0.00 | 5 | 0 | 0.00 | 0 | 0.00 |
| WHO M | 4783 | 35 | 0.73 | 5480 | 0 | 0.00 | 4 | 0 | 0.00 | 0 | 0.00 |
| WHO N | 4772 | 3791 | 79.44 | 5499 | 0 | 0.00 | 4 | 3 | 75.00 | 0 | 0.00 |
| WHO O | 4758 | 7 | 0.15 | 5432 | 0 | 0.00 | 5 | 0 | 0.00 | 0 | 0.00 |
| WHO P | 4630 | 5 | 0.11 | 5277 | 0 | 0.00 | 5 | 0 | 0.00 | 0 | 0.00 |
| WHO U | 4760 | 26 | 0.55 | 5551 | 1 | 0.02 | 5 | 0 | 0.00 | 0 | 0.00 |
| WHO V | 4754 | 2 | 0.04 | 5517 | 0 | 0.00 | 4 | 0 | 0.00 | 0 | 0.00 |
| WHO W | 4893 | 7 | 0.14 | 5613 | 0 | 0.00 | 5 | 0 | 0.00 | 0 | 0.00 |
| WHO X | 4624 | 6 | 0.13 | 5312 | 0 | 0.00 | 4 | 0 | 0.00 | 0 | 0.00 |
| WHO Y | 4765 | 8 | 0.17 | 5510 | 0 | 0.00 | 5 | 0 | 0.00 | 0 | 0.00 |
| WHO Z | 4773 | 12 | 0.25 | 5498 | 0 | 0.00 | 4 | 0 | 0.00 | 0 | 0.00 |

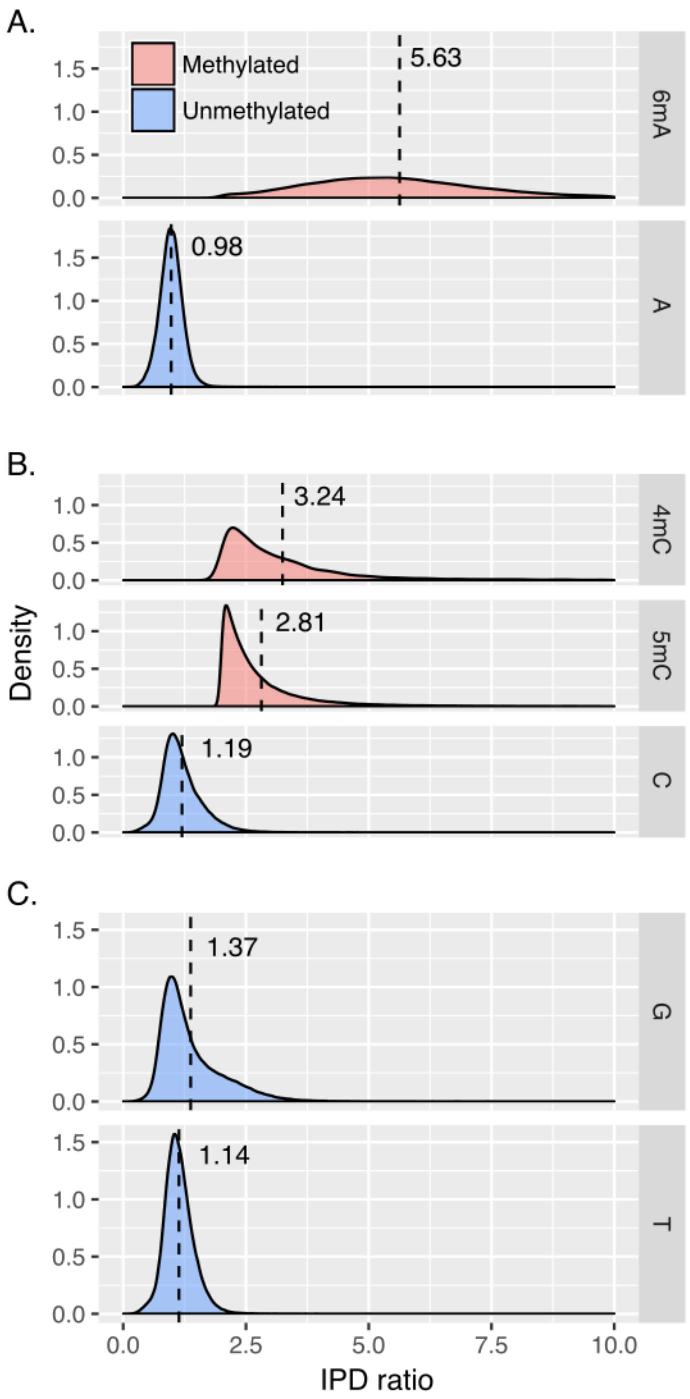


Supplementary Figure 1. Distribution of the IPD ratios for methylated and unmethylated bases. 6mA, 4mC and 5mC IPD values were extracted from the target bases in the final list of curated motifs. Those below an IPD ratio of 2 where excluded to minimize the inclusion of unmethylated motifs. Values for each unmethylated base correspond to a random sample of 10,000 sites per strain outside all the non-redundant predicted motifs detected by the PacBio SMRT pipeline. Dashed vertical lines and numbers indicate the mean value of each distribution.

.

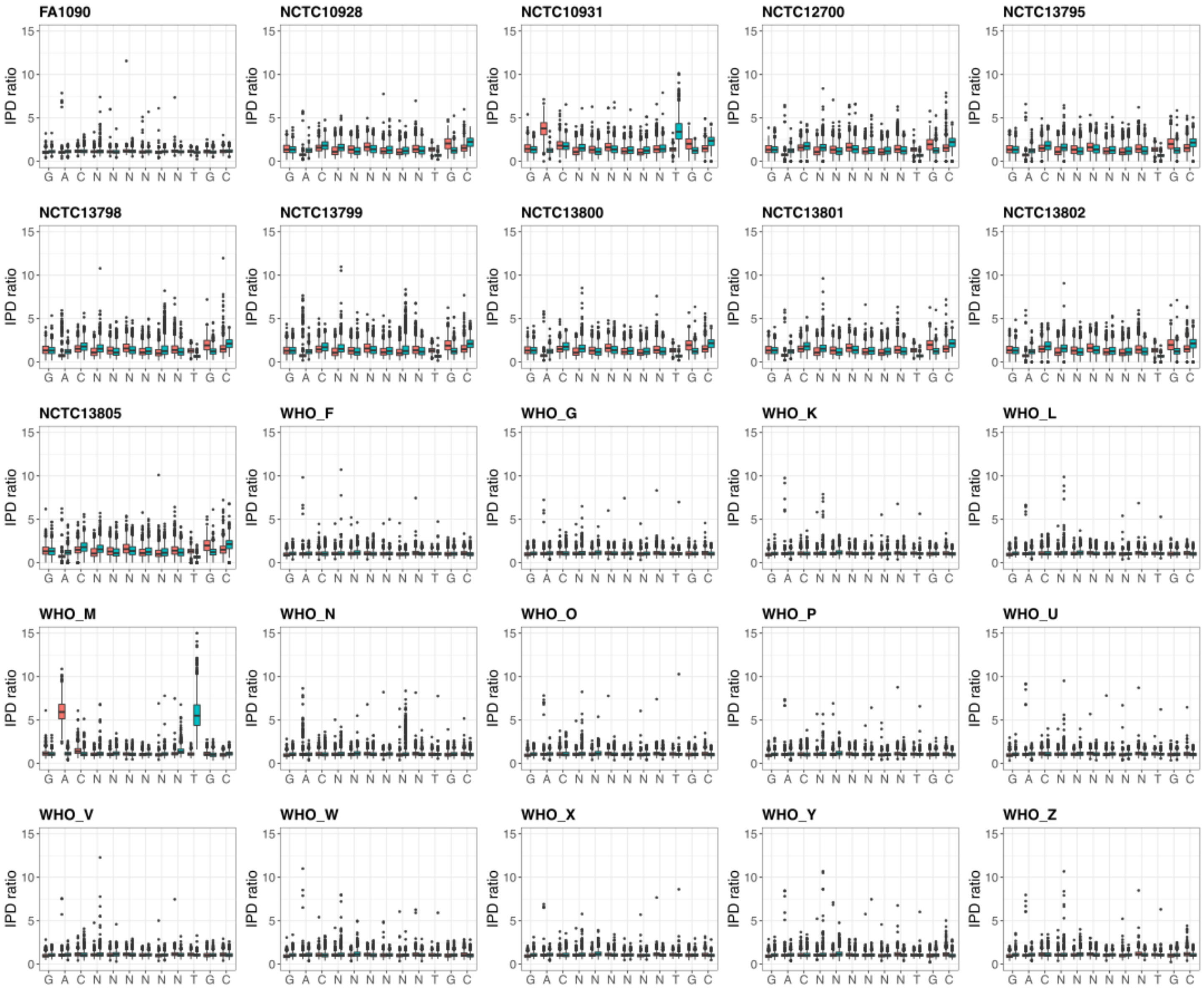


Supplementary Figure 2. IPD ratio values for each base in all instances of the 5’-GACN[6]TGC-3’ motif, target of the Type I NgoAV RMS, in each of the 25 *N. gonorrhoeae* strains included in the study. Per-base distribution of IPD ratio is shown as two separate boxplots containing the values in the forward (red) and reverse (blue) strand.

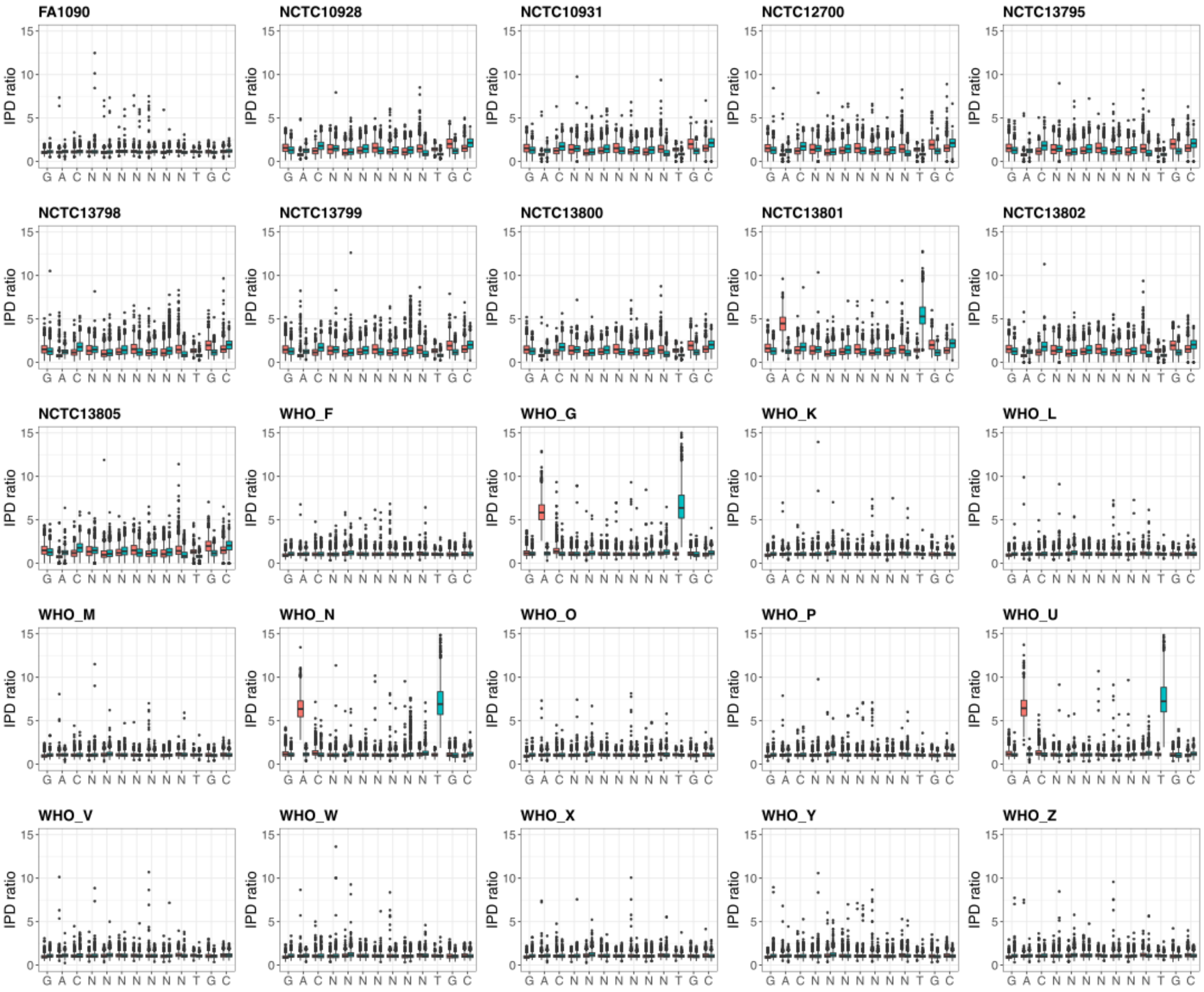


Supplementary Figure 3. IPD ratio values for each base in all instances of the 5'-GACN[7]TGC-3' motif, target of the Type I NgoAV RMS, in each of the 25 *N. gonorrhoeae* strains included in the study. Per-base distribution of IPD ratio is shown as two separate boxplots containing the values in the forward (red) and reverse (blue) strand.

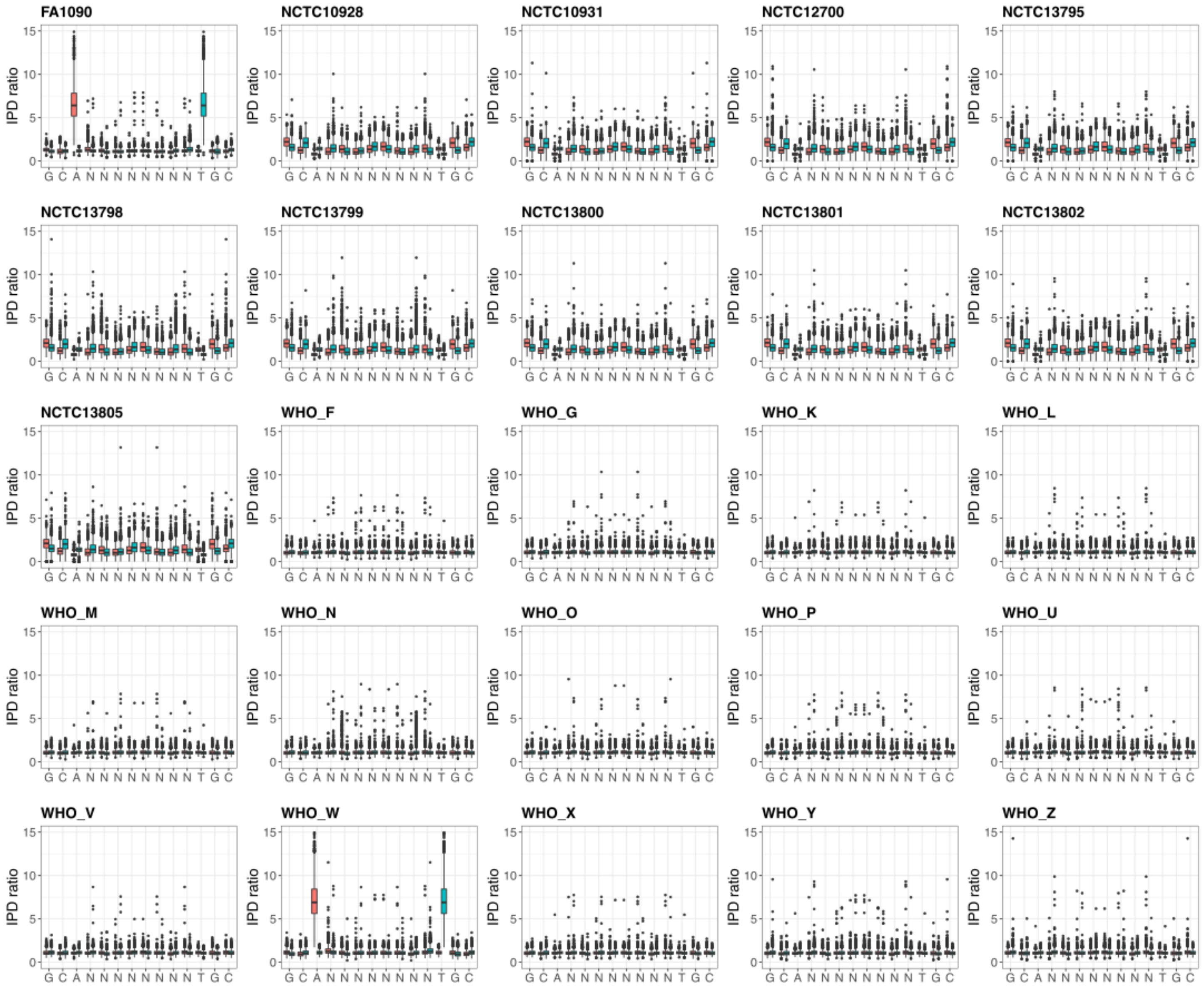


Supplementary Figure 4. IPD ratio values for each base in all instances of the 5'-GCAN[8]TGC-3' motif, target of the Type I NgoAV RMS, in each of the 25 *N. gonorrhoeae* strains included in the study. Per-base distribution of IPD ratio is shown as two separate boxplots containing the values in the forward (red) and reverse (blue) strand.

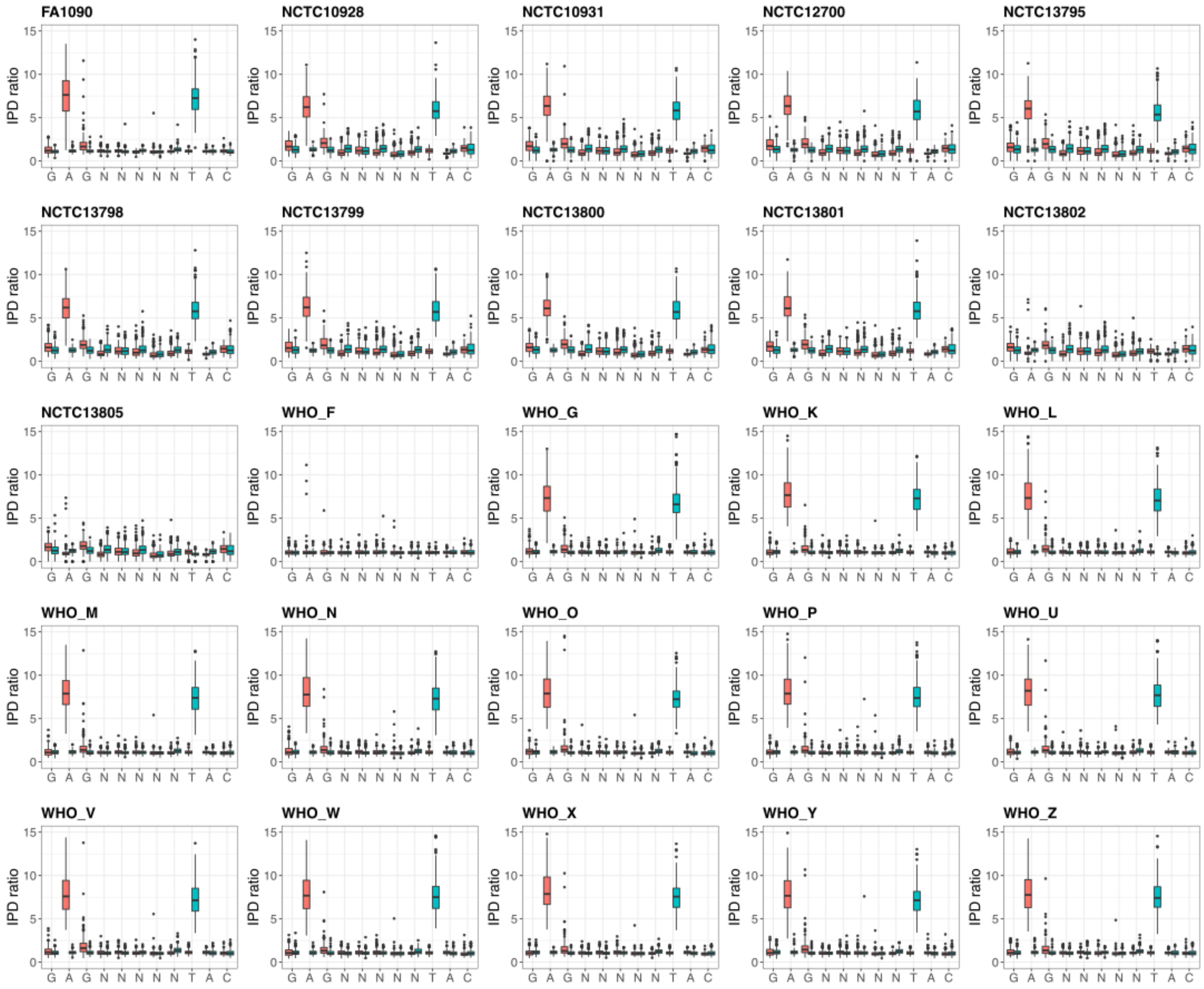


Supplementary Figure 5. IPD ratio values for each base in all instances of the 5'-GAGN[5]TAC-3' motif, target of the Type I NgoAXVIIP RMS, in each of the 25 *N. gonorrhoeae* strains included in the study. Per-base distribution of IPD ratio is shown as two separate boxplots containing the values in the forward (red) and reverse (blue)

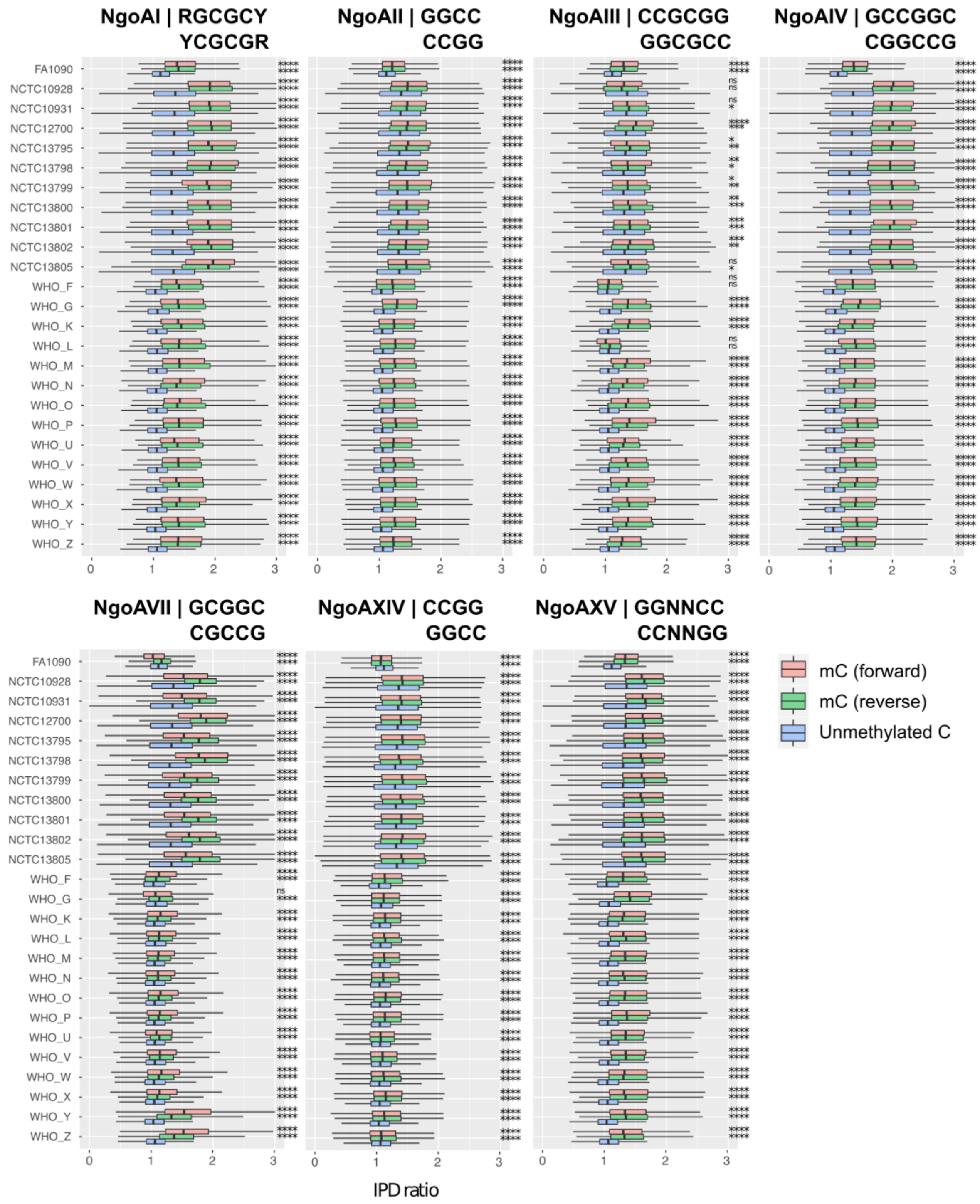


Supplementary Figure 6. IPD ratios for methylated cytosines (mCs) in Type II restriction-modifications system (RMS). The distribution of IPD ratios for the forward (red) and reverse (green) mCs as stated in REBASE are shown as boxplots together with a random 10k subsampling of unmethylated cytosines for each strain (blue). Statistical significance of the comparison among the distribution of the forward and reverse methylations with that from a random 10k subsampling of unmethylated cytosines is indicated next to the boxplots. ****p-value < 0.0001; ***p-value<0.001; **p-value<0.01;*p-value<0.05; ns: non-significant.

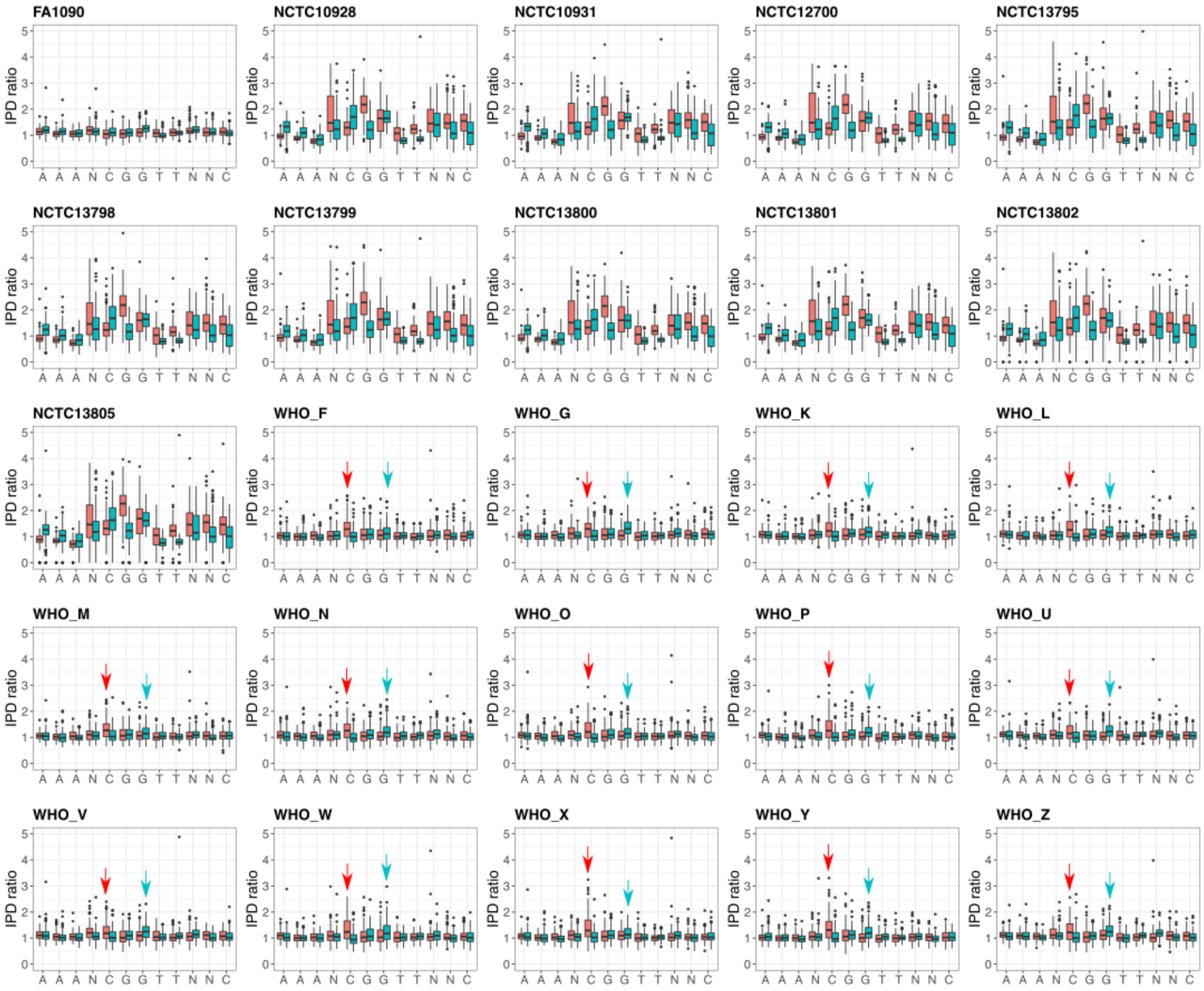


Supplementary Figure 7. IPD ratio values for each base in all instances of the 5'-AAANCGGTTNNC-3' motif in each of the 25 *N. gonorrhoeae* strains included in the study. Per-base distribution of IPD ratio is shown as two separate boxplots containing the values in the forward (red) and reverse (blue) strand.

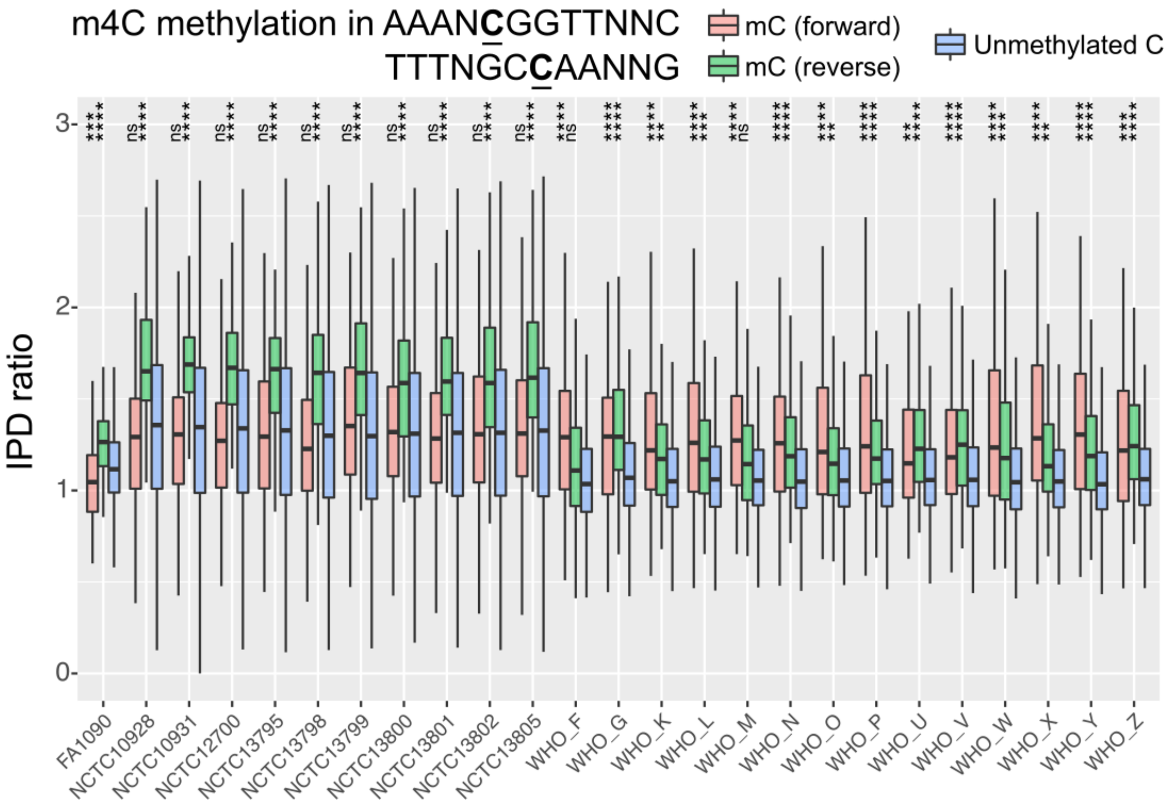


Supplementary Figure 8. IPD ratios for the underlined forward and reverse cytosines in the 5'-AAANCGGTTNNC-3' motif. These values were compared to the distribution of a random 10k subsampling of unmethylated cytosines for each strain. Statistical significance is shown above the boxplots.

****p-value < 0.0001; ***p-value<0.001; **p-value<0.01;*p-value<0.05; ns: non-significant.

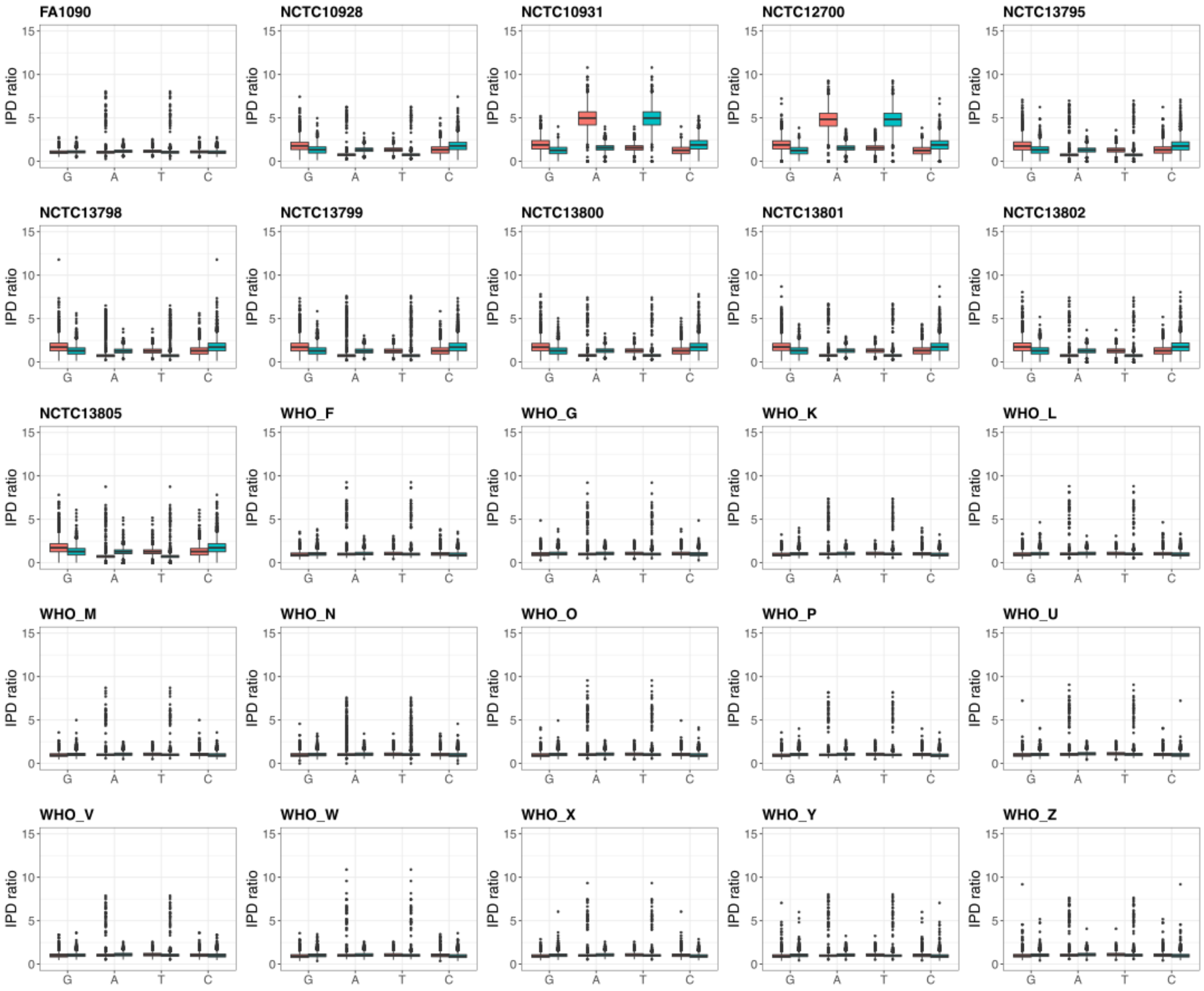


Supplementary Figure 9. IPD ratio values for each base in all instances of the 5'-GATC-3' motif, target of the Type II NgoAXIP RMS, in each of the 25 *N. gonorrhoeae* strains included in the study. Per-base distribution of IPD ratio is shown as two separate boxplots containing the values in the forward (red) and reverse (blue) strand.

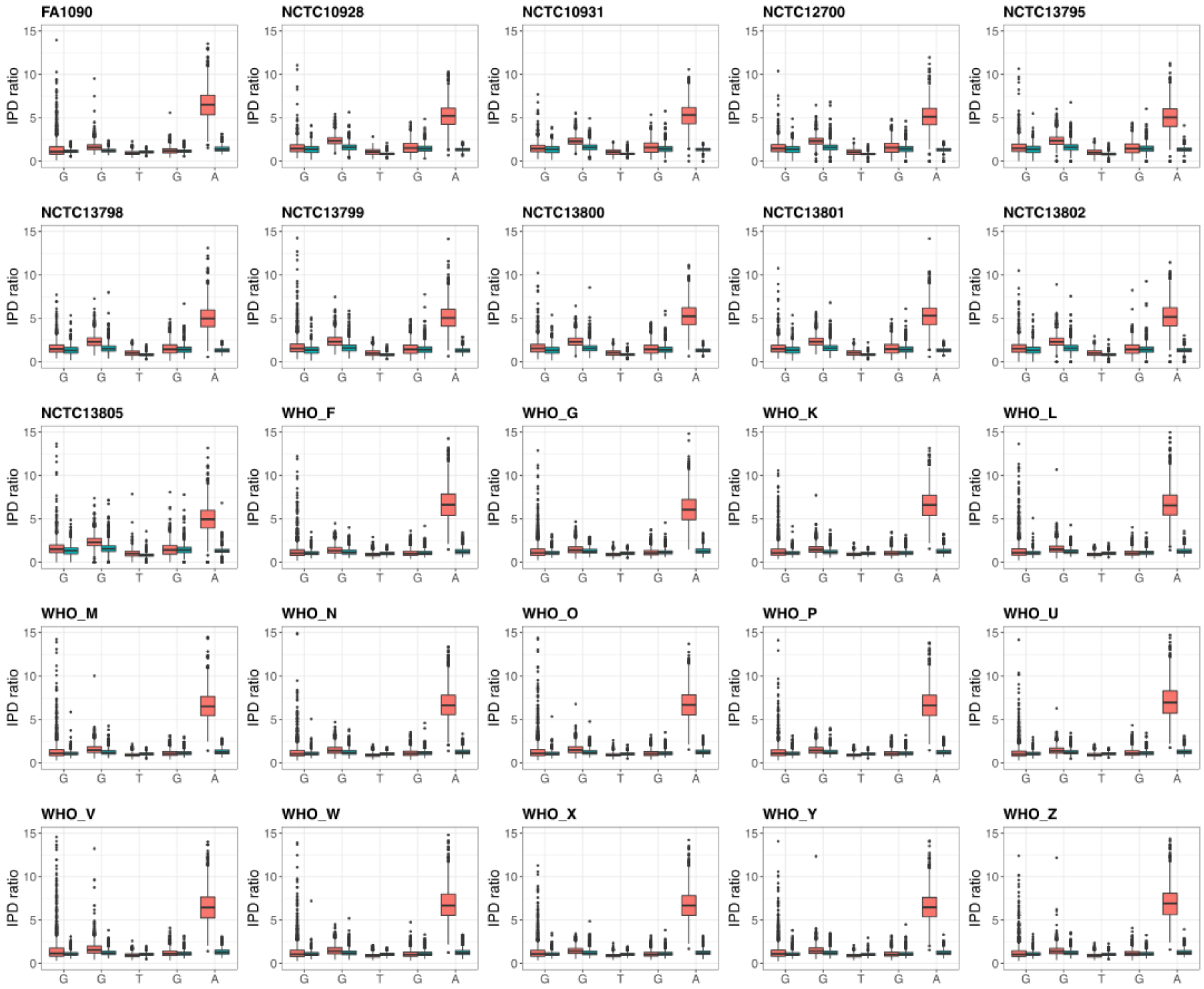


Supplementary Figure 10. IPD ratio values for each base in all instances of the 5'-GGTGA-3' motif, target of the Type II NgoAXVI RMS, in each of the 25 *N. gonorrhoeae* strains included in the study. Per-base distribution of IPD ratio is shown as two separate boxplots containing the values in the forward (red) and reverse (blue) strand.

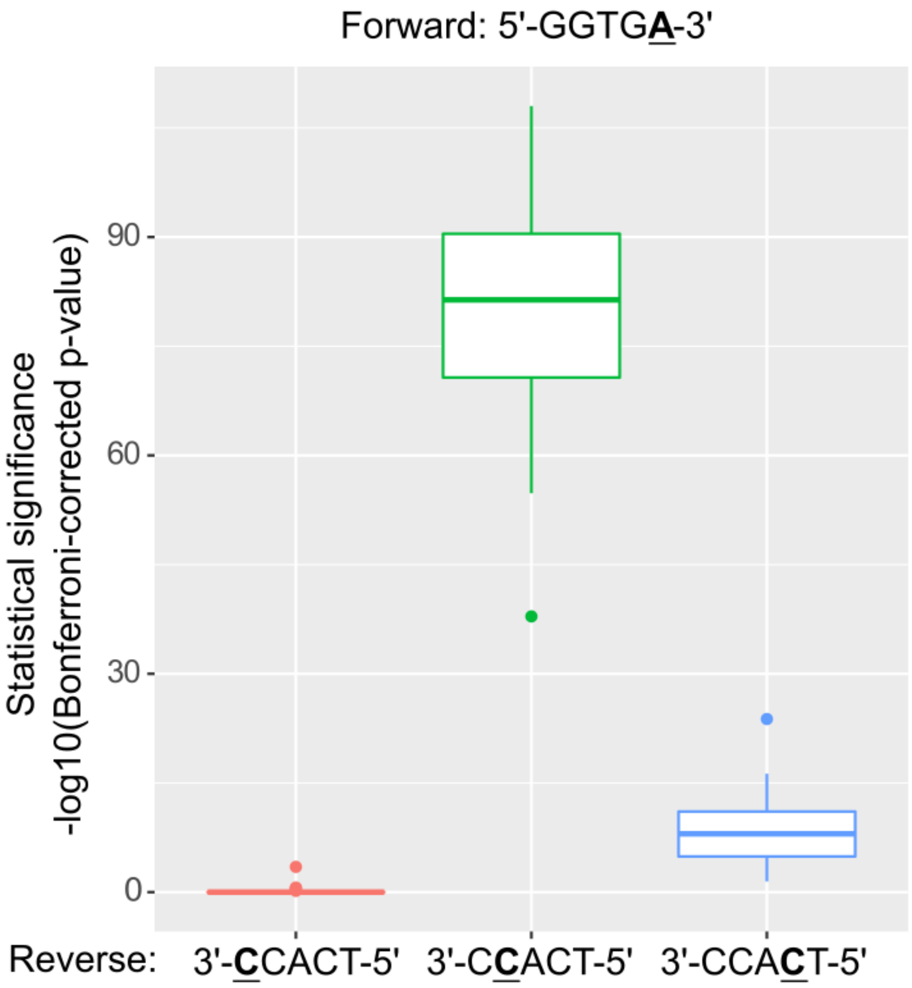


Supplementary Figure 11. Inference of the methylated cytosine in the reverse strand of 5'-GGTGA-3' (NgoAXVI). Bonferroni-corrected p-values are shown in logarithmic scale for the comparison between the distribution of IPD values for the underlined cytosine and a random 10k subsampling of unmethylated cytosines for each strain. The cytosine with a higher significance and thus, candidate to be methylated, is the second one.

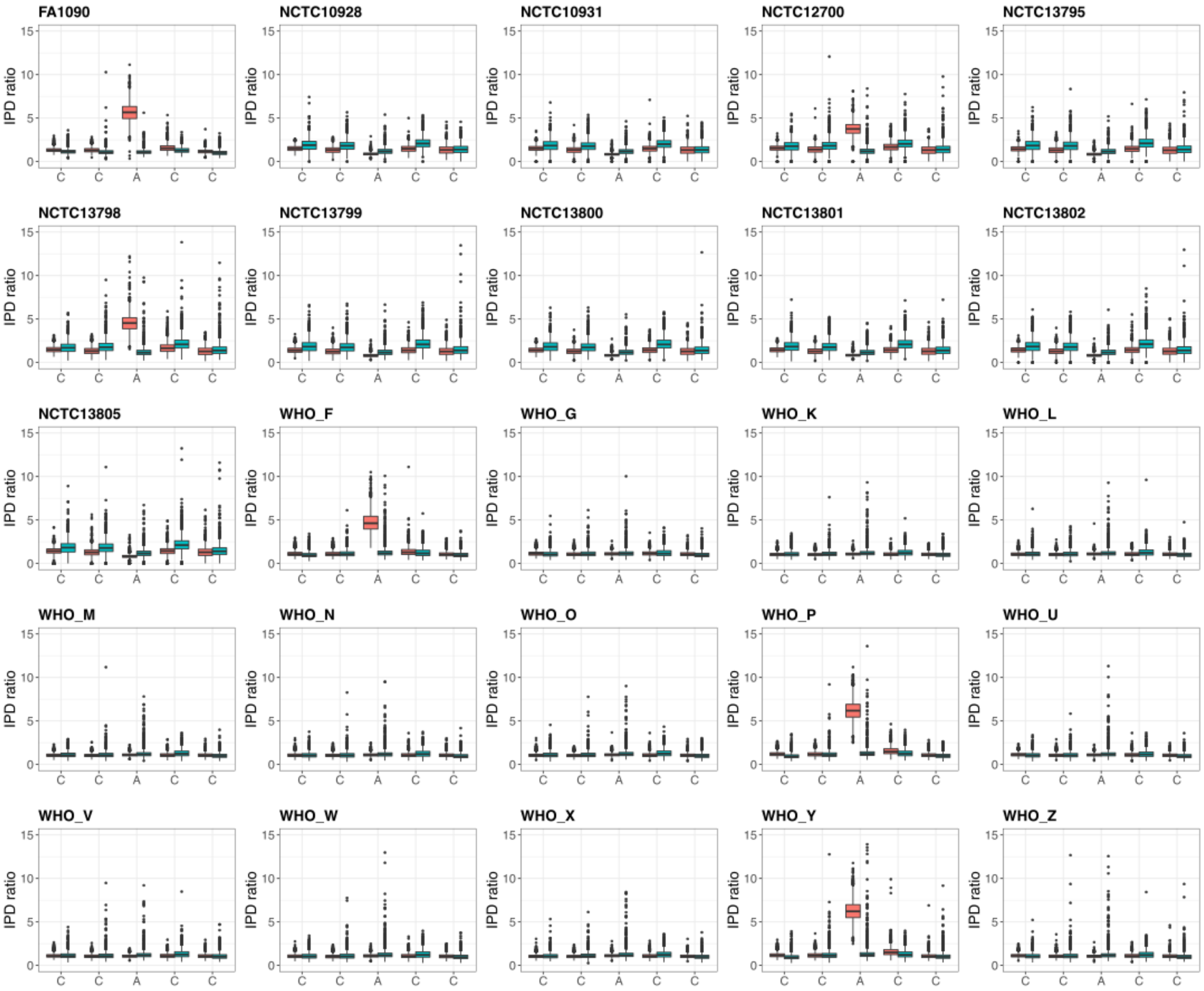


Supplementary Figure 12. IPD ratio values for each base in all instances of the 5'-CCACC-3' motif, target of the Type III NgoAX RMS, in each of the 25 *N. gonorrhoeae* strains included in the study. Per-base distribution of IPD ratio is shown as two separate boxplots containing the values in the forward (red) and reverse (blue) strand.

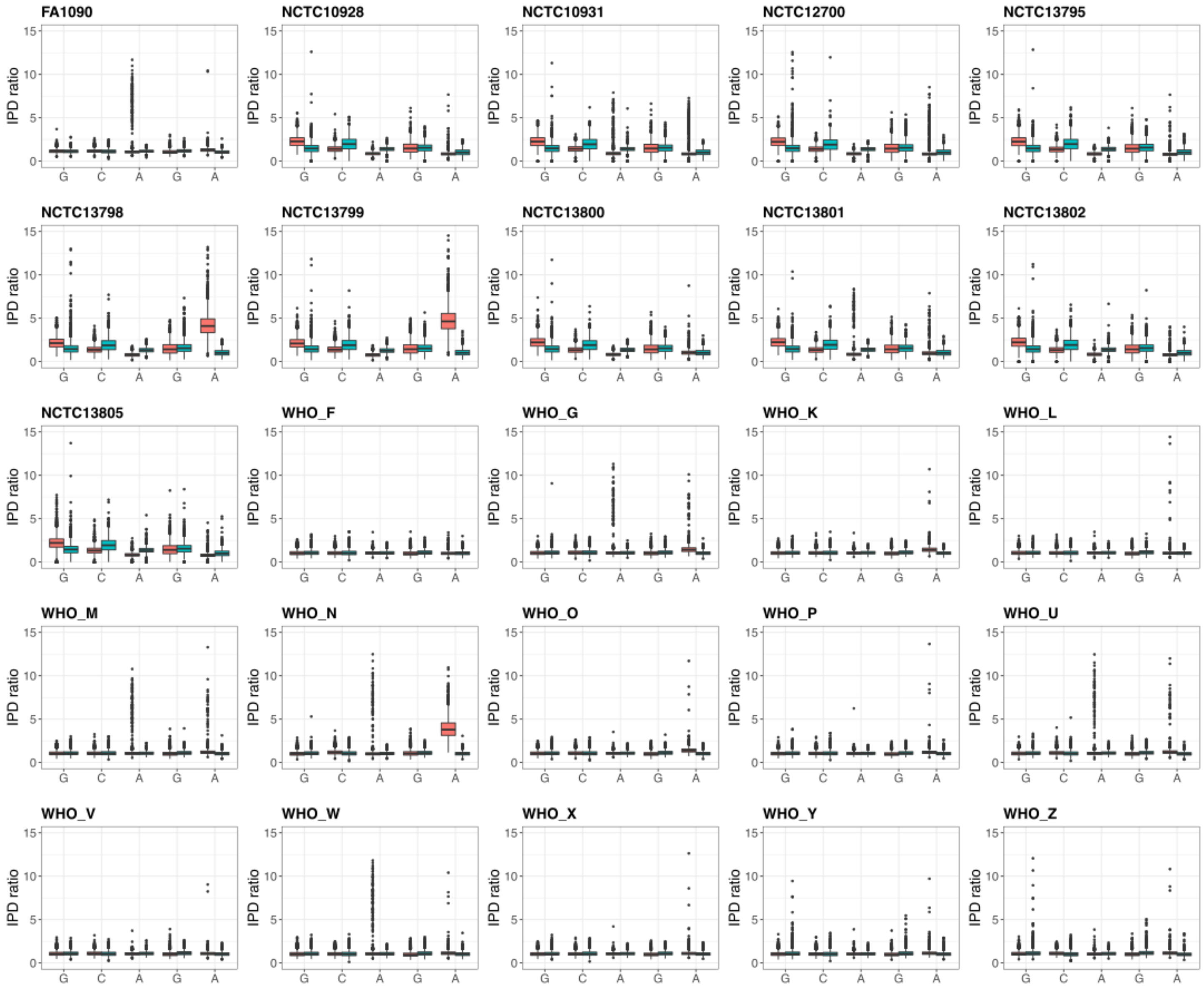


Supplementary Figure 13. IPD ratio values for each base in all instances of the 5'-GCAGA-3' motif, target of the Type III NgoAXIIP RMS, in each of the 25 *N. gonorrhoeae* strains included in the study. Per-base distribution of IPD ratio is shown as two separate boxplots containing the values in the forward (red) and reverse (blue) strand.

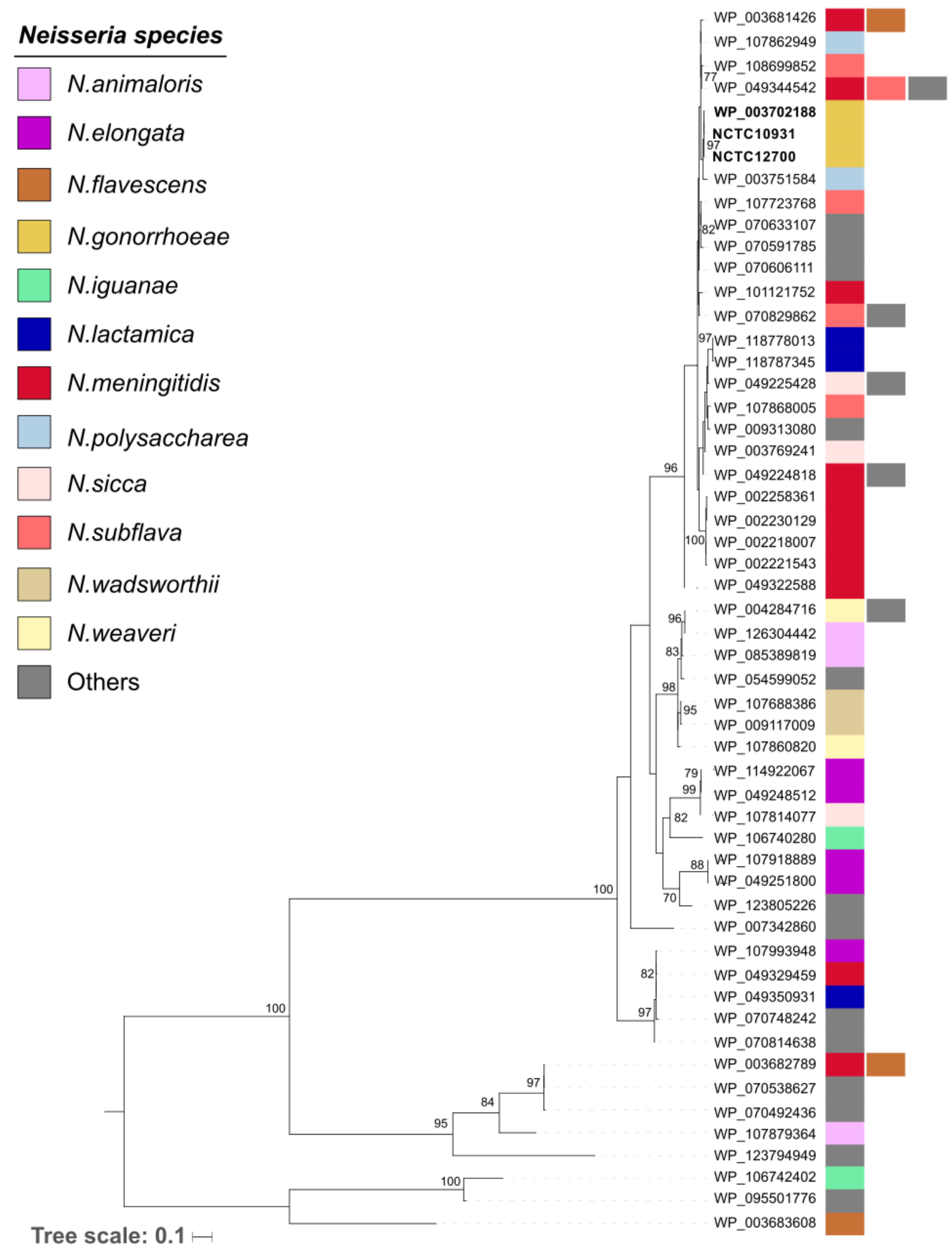


Supplementary Figure 14. Maximum likelihood phylogenetic reconstruction of representative Dam protein sequences available in genomes of the *Neisseria* genus in the RefSeq database. These representative sequences (WP_*) were obtained using the Identical Protein Groups (IPG) tool in NCBI. Coloured strips represent the species in which these sequences are found. Those appearing only once have been coloured in grey as 'Others' but can be accessed using the ID on the tip labels in [https://www.ncbi.nlm.nih.gov/ipg](https://www.ncbi.nlm.nih.gov/ipg).

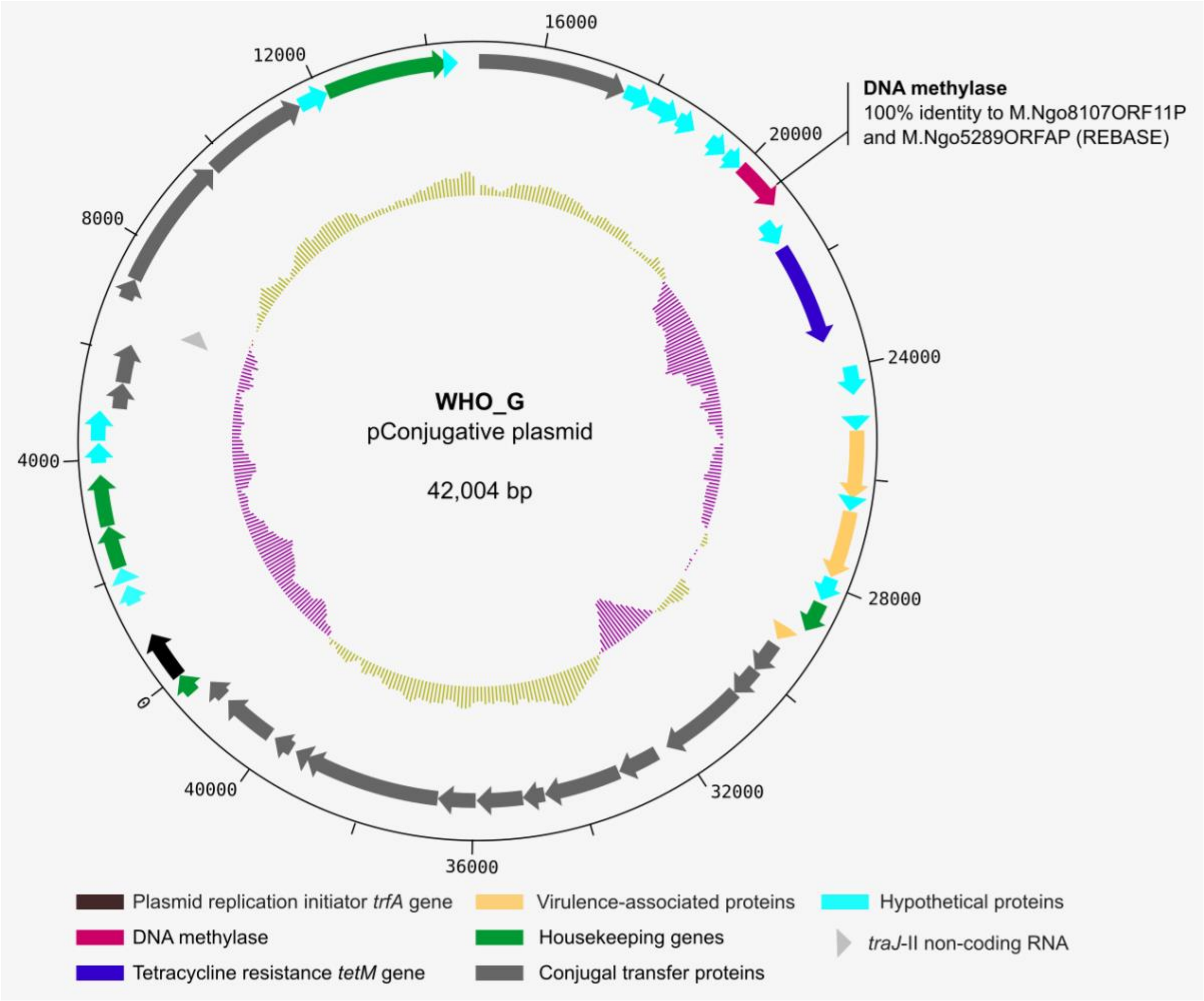


Supplementary Figure 15. Circular plot (58) representing the annotation of the WHO_G pConjugative plasmid, which shows the presence of an orphan DNA methylase that has a 100% protein identity with two entries in REBASE.
